# Characterizing the differential distribution and targets of Sumo paralogs in the mouse brain

**DOI:** 10.1101/2022.09.09.507035

**Authors:** Terry R. Suk, Trina T. Nguyen, Zoe A. Fisk, Miso Mitkovski, Haley M. Geertsma, Jean-Louis A. Parmasad, Meghan M. Heer, Steve M. Callaghan, Nils Brose, Marilyn Tirard, Maxime W.C. Rousseaux

## Abstract

SUMOylation is an evolutionarily conserved and essential mechanism whereby Small Ubiquitin Like Modifiers, or SUMO proteins (Sumo in mice), are covalently bound to protein substrates in a highly dynamic and reversible manner. SUMOylation is involved in a variety of basic neurological processes including learning and memory, and central nervous system development, but is also linked with neurological disorders. However, studying SUMOylation *in vivo* remains challenging due to limited tools to study Sumo proteins and their targets in their native context. More complexity arises from the fact that Sumo1 and Sumo2 are ∼50% homologous, whereas Sumo2 and Sumo3 are nearly identical and indistinguishable with antibodies. While Sumo paralogues can compensate for one another’s loss, Sumo2 is highest expressed and only paralog essential for embryonic development making it critical to uncover roles specific to Sumo2 *in vivo*. To further examine the roles of Sumo2, and to begin to tease apart the redundancy and similarity between key Sumo paralogs, we generated (His_6_-)HA epitope-tagged Sumo2 knock-in mouse alleles, expanding the current Sumo knock-in mouse tool-kit comprising of the previously generated His_6_-HA-Sumo1 knock-in model. Using these HA-Sumo mouse lines, we performed whole brain imaging and mapping to the Allen Brain Atlas to analyze the relative distribution of the Sumo1 and Sumo2 paralogues in the adult mouse brain. We observed differential staining patterns between Sumo1 and Sumo2, including a partial localization of Sumo2 in nerve cell synapses of the hippocampus. Combining immunoprecipitation with mass spectrometry, we identified native substrates targeted by Sumo1 or Sumo2 in the mouse brain. We validated select hits using proximity ligation assays, further providing insight into the subcellular distribution of neuronal Sumo2-conjugates. These mouse models thus serve as valuable tools to study the cellular and biochemical roles of SUMOylation in the central nervous system.

## INTRODUCTION

Post translational modifications (PTMs) provide a dynamic mode of regulation over protein functions altering the secondary, tertiary, or quaternary structures of proteins. Although the most frequently characterized PTMs typically involve the covalent conjugation of small molecules, such as phosphate, glycosyl, or acetyl groups, protein function can also be modulated by the covalent conjugation of large molecules such as the protein Ubiquitin. Indeed, several Ubiquitin-like proteins have been identified to function as PTMs in eukaryotes including NEDD8, Atg8, Atg12, and Small Ubiquitin-like Modifiers (SUMOs) (Ilic et al., 2022; Mahajan et al., 1997; Matunis et al., 1996).

The covalent addition of SUMOs through SUMOylation is a dynamic and essential process that is highly conserved throughout eukaryote evolution(Celen and Sahin, 2020; Ilic et al., 2022). First, immature SUMO proteins are matured through Sentrin Specific Protease (SENP) cleavage at a diglycine motif that is essential for covalent conjugation (Xu and Au, 2005). Next, SUMOs are activated in an ATP dependent manner and conjugated to the E1 ligase heterodimer complex consisting of SUMO Activating Enzyme 1 (SAE1) and Ubiquitin-Like Modifier Activating Enzyme 2 (UBA2)(Gong et al., 1999). SUMOs are then transferred to the sole E2 ligase, SUMO conjugating enzyme UBC9 (UBC9, encoded by *UBE2I*), which is essential for SUMOylation to occur(Desterro et al., 1997; Gong et al., 1997; Johnson and Blobel, 1997; Lee et al., 1998; Saitoh et al., 1998; Seufert et al., 1995). SUMOylation can then occur at a lysine residue of a target substrate, typically residing in a SUMO consensus motif (Ψ-K-x-D/E, Ψ=large hydrophobic amino acid), directly from UBC9 or upon facilitation by E3 ligases (Rodriguez et al., 2001; Sampson et al., 2001). Finally, SENPs can readily deSUMOylate the protein substrate, and SUMO is recycled for subsequent rounds of SUMOylation, allowing for highly dynamic regulation of substrate function (Nayak and Müller, 2014).

There are five known human SUMO paralogs (SUMO1 – SUMO5) with SUMO1 – SUMO3 being the most extensively studied as they are ubiquitously expressed and found in most vertebrates, whereas SUMO4 and SUMO5 are specific to humans and exhibit tissue-specific expression (Bouchard et al., 2021; Citro and Chiocca, 2013). SUMO1 shares around 50% homology with SUMO2 and SUMO3, whereas mature SUMO2 and SUMO3 share 97% homology leading to these proteins to be often referred together as SUMO2/3 (Bohren et al., 2004). While SUMO paralogs often play redundant roles and can exhibit some level of compensation, each paralog can also play unique roles in cells, localizing to different subcellular regions and targeting different protein substrates (Citro and Chiocca, 2013; Ilic et al., 2022). Additionally, *Sumo2*, but not *Sumo1* nor *Sumo3*, is the only essential paralog whereby knock-out in mice results in major developmental defects leading to embryonic lethality by E10.5 (Wang et al., 2014b). The essentiality of Sumo2 for proper development may be in part due to higher levels of Sumo2 in comparison to its paralogs, particularly Sumo3 (Wang et al., 2014b; Yu et al., 2020). Alternatively, it may also be the result of specific Sumo-conjugated proteomes between Sumo2 and its paralogs conferring different roles in cells. Thus, characterizing the specific roles of Sumo2 is crucial to not only help to uncover key mechanisms of cellular function and advance our understanding of SUMOylation in basic biology, but to also provide insight into disease processes.

Several challenges currently limit the study of protein SUMOylation: 1) Differentiating between the Sumo paralogs using antibodies is limiting or not possible due to the high degree of homology(Garvin et al., 2022); 2) The highly dynamic nature of SUMOylation makes biological context such as tissue, cell type, and cell state, critical for capturing and characterizing SUMOylation events; and 3) Tools to study SUMOylation are limited and often rely on overexpression models, potentially confounding studies of protein function. Consequently, protein SUMOylation is heavily studied in highly proliferating cells where it is now well established that SUMOylation plays a key role in regulating nuclear and DNA-related mechanisms, but the role of this PTM remains unclear, and even debated, in more complex system, including post-mitotic cells such as neurons (Daniel et al., 2018; Daniel et al., 2017).

To overcome these limitations, we expanded the current Sumo mouse knock-in toolkit and generated and characterized in parallel two novel knock-in (KI) mouse models. Specifically, sequences encoding affinity tags are knocked into the endogenous *Sumo2* locus whereby an HA-Sumo2 or His_6_-HA-Sumo2 fusion protein is generated, allowing for the characterization of Sumo2 and its substrates *in vivo*. Analogous to our work with a corresponding His_6_-HA-Sumo1 KI allele (Daniel et al., 2018; Daniel et al., 2017; Tirard et al., 2012), this approach allows us to directly compare the localization patterns between Sumo1 and Sumo2 throughout the mouse central nervous system (CNS). We used whole brain imaging to map the relative abundance of Sumo1 and Sumo2 throughout the adult mouse brain and observed that, while Sumo levels are broadly distributed, they exhibit regional differences throughout the CNS. We found that, unlike Sumo1 (Daniel et al., 2018; Daniel et al., 2017; Tirard et al., 2012), Sumo2 is present in both nuclear and extra-nuclear compartments, including neuronal synapses. Based on this distinct subcellular distribution, we explored native Sumo1 and Sumo2 targets using immunoprecipitation and mass spectrometry. We identified genuine Sumo2-specific targets in the adult mouse brain, including a subset of non-nuclear proteins. Moreover, we observed dynamic changes between Sumo1- vs. Sumo2-selective substrates. Finally, we validated several hits using proximity ligations assays to detect protein-Sumo interactions within primary cortical neurons. This approach did not only provide an additional mode of validating native interactors of Sumo2 in wild-type neurons, but also yielded spatial information, demonstrating both nuclear and cytoplasmic interactions between Sumo2 and its substrates. Together, these new mouse lines and our data provide an important new resource that lays the foundation of a “Sumo-code” which provides a new layer of complexity to brain function.

## RESULTS

### Generation of (His_6_-)HA-Sumo2 mouse lines

The use of epitope tags provides a versatile way to streamline the study of proteins by taking advantage of the highly selective nature of antibodies raised against these tags. In parallel institutions, we generated two independent alleles to facilitate the study of Sumo2 *in vivo*. We took advantage of a CRISPR/Cas9 approach to knock-in a 6xHis-hemagglutinin (*His*_*6*_*-HA*) tag into the amino-terminus of *Sumo2* (*His*_*6*_-*HA-Sumo2*) in C57BL/6J mice (Figure 1—figure supplement 1A) in one case, mirroring that of a previously generated mouse line to study Sumo1 *in vivo* (Tirard et al., 2012). Moreover, we also generated an HA-Sumo2 knock-in mouse that does not have the His_6_ tag. We observed that knock-in of either *His*_*6*_*-HA* or *HA* to *Sumo2* resulted in hydrocephaly and premature death in heterozygous mice, indicating a likely hypomorphic effect of the epitope tag on a C57BL/6J background. Interestingly, backcrossing the mice to an FVB/N background alleviated these effects, resulting in no overt differences between mutant and wild-type FVB/N (WT) littermates in either line. Nevertheless, given the importance of Sumo2 for life, and the robustness of the HA tag for visualization and biochemistry, we performed the bulk of our analyses on heterozygous His_6_-HA-Sumo2 (and heterozygous His_6_-HA-Sumo1; also crossed to an FVB/N background) knock-in mice.

### His_6_-HA-Sumo2 enables the comparison of endogenous Sumo2 levels in the CNS

To determine whether the addition of the HA epitope tag affects native Sumo levels, we assessed the levels of Sumo2/3 conjugates by Western blot, comparing total levels of unconjugated Sumo (“Free Sumo”) and Sumo-conjugated proteins from mouse brain lysates. We observed no change in the abundance of Sumo2/3 conjugates in the brains of adult His_6_-HA-Sumo2 mice compared to His_6_*-*HA-Sumo1 or WT mice (Figure 1—figure supplement 1B). As decreased levels of Sumo1 conjugates were previously observed in homozygotes His_6_-HA-Sumo1 mouse brain (Daniel et al., 2017), we thus measured whether there were any changes in the abundance of native Sumo1 conjugates in heterozygous His_6_-HA-Sumo1 and His_6_-HA-Sumo2 KI mice (Wang et al., 2014b). Again, no changes were observed between either Sumo KI line and their WT counterparts, indicating that the heterozygous His_6_-HA-Sumo2 or His_6_-HA-Sumo1 allele does not alter overall levels of Sumo conjugates in mouse brain (Figure 1—figure supplement 1C). Additionally, we did not observe a significant effect on the expression of *Sumo1* and *Sumo2* transcript levels in either of the His_6_-HA-Sumo KI mice (Figure 1––figure supplement 1D). Thus, the novel His_6_-HA-Sumo2 KI mice model enable the study of Sumo2 at endogenous levels *in vivo* without changes to the Sumo equilibrium.

The uniformity of the HA-tag epitope between the His_6_-HA-Sumo1 and His_6_-HA-Sumo2 mice allows for a more direct comparison of endogenous Sumo1 and Sumo2 levels *in vivo*. We assessed the relative levels of Sumo1 and Sumo2 in the central nervous system, specifically the cortex (CTX), cerebellum (CBM), olfactory bulbs (OB), spinal cord (SC) and remainder of the brain (termed “striatum-thalamus-brainstem for simplicity [S-Th-B]”) from adult mice (Figure 1). Western blot analysis from CNS regional lysates shows that levels of unconjugated and conjugated Sumo2 are significantly higher than Sumo1 throughout the CNS (2-way ANOVA; Sumo effect: *P* = 0.0046 and *P* = 0.0003 for high molecular weight Sumo and free Sumo, respectively). The CTX, CBM, and OB are specifically enriched for Sumo2 whereas the S-Th-B and SC contains lower levels of Sumo2. Taken together, Sumo1 and Sumo2 are broadly distributed throughout the CNS, however regional differences in Sumo abundance and Sumo paralog predominance can be observed, indicating possible unique regional roles throughout the CNS.

**Figure 1:**
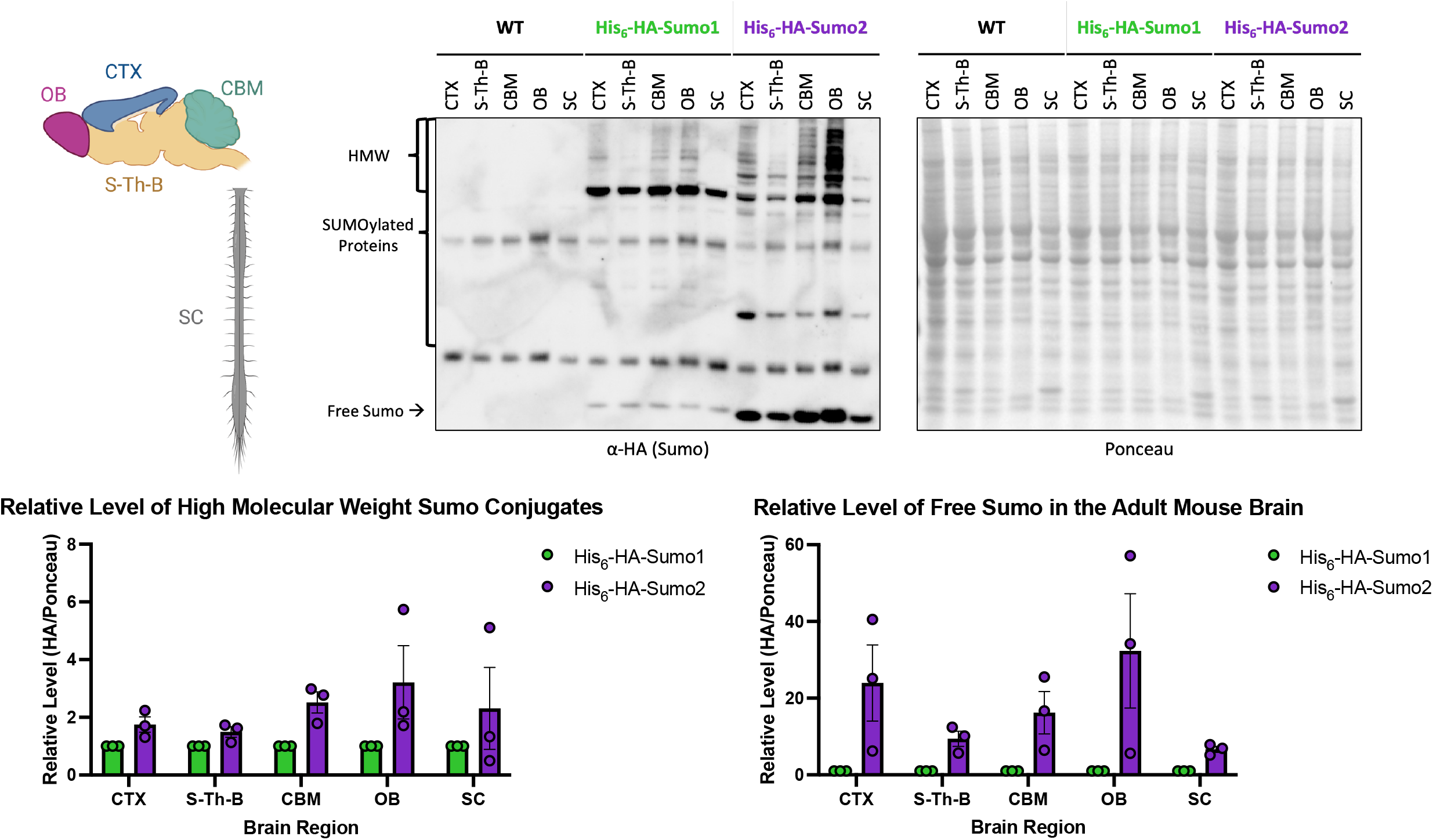
HA-epitope knock-in enables direct comparisons of differential levels of Sumo1 and Sumo2 and their conjugates in the murine central nervous system. Western blot analysis of various regions of His_6_-HA-Sumo1 and His_6_-HA-Sumo2 mouse central nervous system. Brain regions highlighted in the schematic (Top Left) were dissected for protein extraction and analyzed by Western blot using anti-HA antibody. HA signal corresponding to SUMOylated protein (bracket on the left side of the top left panel, HMW: High Molecular Weight) was normalized to ponceau (right top panel) and quantified via densitometry (bottom panels), N=3.

### Regional Analysis of Sumo1 and Sumo2 Throughout the Mouse Brain

As differences in Sumo levels are detectable from regional crude brain lysates, we leveraged the specificity of the HA-tag and performed whole brain clearing and mapping of Sumo1 and Sumo2 signals in the adult mouse brain. His_6_-HA-Sumo1 and His_6_-HA-Sumo2 mouse brains were cleared and immunolabeled against the HA-tag and NeuN before imaging using light sheet microscopy (Figure 2A; Figure 2—figure supplement videos 1 & 2). Images were then analyzed and aligned to the Allen Brain Atlas to determine the relative levels of the His_6_-HA-Sumo signals in defined brain regions (Table 1). It is worth noting that due to the different absolute abundance of Sumo1 (low) versus Sumo2 (high) in the brain, only qualitative Sumo1 vs Sumo2 comparisons were made. To generate relative maps of HA-Sumo distribution, the mean regional HA signal intensity was first normalized to the regional density of NeuN to account for changes in cell density before being averaged across hemibrains to account for tissue variability. Generally, Sumo1 and Sumo2 showed similar distributions throughout the mouse brain (Figure 2B-E) and we observed that both Sumo1 and Sumo2 were most abundant in cerebellar nuclei.

**Figure 2:**
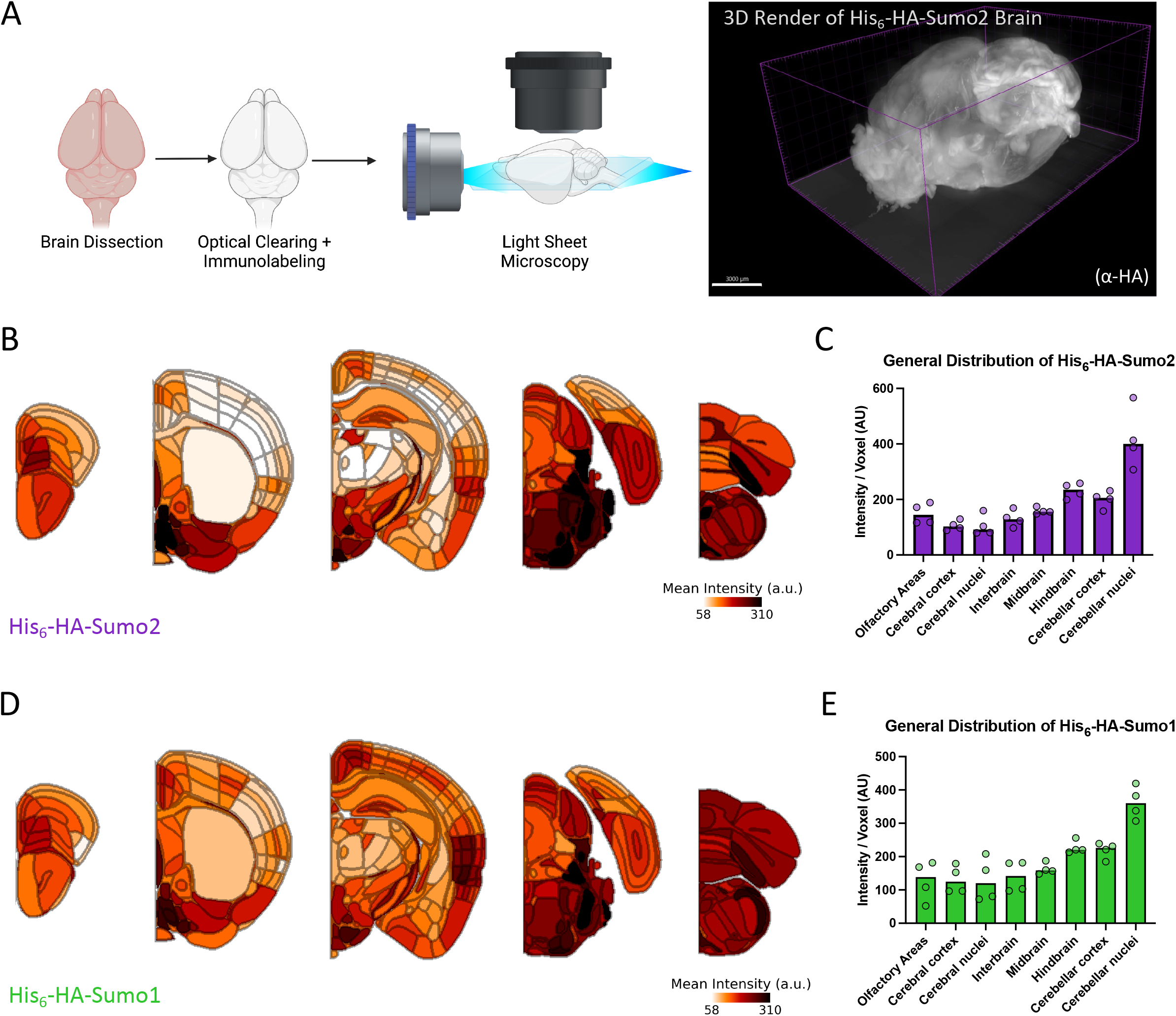
Whole brain imaging reveals the topographical distribution of Sumo paralogs. (A). Schematic of brain clearing and light sheet microscopy for whole brain imaging with a 3D Imaris render of a His_6_-HA-Sumo2 brain. (B). Heatmaps of anti-HA immunosignal intensity normalized to regional density quantified by NeuN staining and alignment to the Allen Brain Atlas across the His_6_-HA-Sumo2 KI depicted brain region, averaged between hemibrains and biological replicates (N=2, two hemispheres). (C). Bar plot depicting the relative levels of anti-HA immunosignal per voxel averaged across across His_6_-HA-Sumo2 brain regions defined by the Allen Brain Atlas. (D). Heatmaps of anti-HA immunosignal intensity normalized to regional density quantified by NeuN staining and alignment to the Allen Brain Atlas across the His_6_-HA-Sumo1 KI depicted brain region, averaged between hemibrains and biological replicates (N=2, two hemispheres). (E). Bar plot depicting the relative levels of anti-HA immunosignal intensity normalized to regional density across His_6_-HA-Sumo1 brain regions defined by the Allen Brain Atlas. Each datapoint in (C) and (D) is from the mean intensity from a single hemisphere (N=2, two hemispheres).

While Sumo levels were generally evenly distributed when characterizing gross anatomical regions, clear nuances in Sumo abundance were detectable at higher anatomical resolution. For example, in the hippocampus, Sumo levels were highest in the parasubiculum and presubiculum, and lowest in the *Fasciola cinerea* (Figure 2—figure supplement 1A & B). In the neocortex, we observed that Sumo levels were generally enriched in layers 2/3 (Figure 2B & D; Figure 2—figure supplement 1C), an area critical for integrative processing (Feldmeyer, 2012). Interestingly, while Sumo1 and Sumo2 often share similar distributions, we observed that Sumo2 is enriched in the anterior hypothalamus (Figure 2—figure supplement 1D), particularly in regions involved in the circadian rhythm, including the suprachiasmatic nucleus and subparaventricular zone (Drunen and Eckel-Mahan, 2021).

We investigated whether the brain-wide imaging findings could be validated by an orthogonal immunofluorescence assay (Figure 2—figure supplement 2). Looking at multiple brain regions, we found that Sumo1 and Sumo2 are broadly distributed through the somatosensory cortex. Interestingly, Sumo1 maintains a strictly nuclear localization, whereas Sumo2 is observed throughout the nucleus and in the cytoplasm and axons of cortical neurons. In the cerebellum, we found that Sumo2 is specifically enriched in granule cells, contrasting the strong Purkinje cell staining observed in His_6_-HA-Sumo1 mice (Tirard *et al*. 2012). The hippocampus also displays characteristic nuclear staining for His_6_-HA-Sumo1 in both the CA1 and CA3, whereas Sumo2 is seen throughout the nucleus and in the somata, and axons of hippocampal neurons. Thus, while Sumo1 and Sumo2 may share regional expression in the brain, the distinct localizations of Sumo1 and Sumo2 *in vivo* indicate different targets of SUMOylation or different metabolism or conjugation dynamics of free Sumo likely play unique roles in neuronal regulation.

### HA-Sumo2 localizes to synapses in vivo

Consistent with our previous work, we observed that Sumo1 strictly localizes to neuronal nuclei (Figure 2–figure supplement 2) (Tirard et al., 2012). However, we noticed that Sumo2 signal can also be found outside the nuclear compartment (Figure 2–figure supplement 2). The role of Sumo in extra-nuclear compartments, specifically at synapses, remains a topic of debate, particularly in the context of Sumo1(Daniel et al., 2018; Daniel et al., 2017). Thus, we further characterized extra-nuclear Sumo2 signals using homozygous HA-Sumo2 mice. As in the His_6_-HA-Sumo2 KI model, levels of Sumo2 conjugates were unaltered in total brain lysates from homozygous HA-Sumo2 mice as compared to WT littermates, ruling out an effect of the HA tag on overall Sumo2 expression and conjugation levels (Figure 3–figure supplement 1A) (Tirard et al., 2012). Western blot analysis of brain subcellular fractions using an anti-HA antibody highlighted not only the difference in global levels of Sumo1 and Sumo2 conjugates, but also revealed that Sumo2 conjugates are abundant in non-nuclear fractions, including synaptic cytosol fractions S2 and S3 (Figure 3–figure supplement 1B-C). A weak but specific signal was also observed in crude synaptosomes and other synaptic membrane fractions (Figure 3– figure supplement 1B-D).

Next, we used homozygous HA-Sumo2 mice to further characterize extra-nuclear Sumo2 by performing anti-HA immunolabeling of the hippocampal CA3 area, a region enriched in synapses (Figure 2–figure supplement 2). Brain sections from WT and HA-Sumo2 mice were immunolabeled for HA and neuronal markers for dendrites and synaptic compartments, and anti-HA immunosignals were quantified in HA-Sumo2 mice as compared to WTs (Figure 3). Using confocal microscopy, we confirmed the presence of Sumo2 in extra-nuclear compartments, specifically along dendrites and in synapses (Figure 3A). Quantification of HA-Sumo2 immunosignals in DAPI positive area confirmed the strong enrichment of Sumo2 conjugates in neuronal nuclei, in agreement with the well-known role of Sumos as nuclear regulators (Figure 3A and B) (Hendriks and Vertegaal 2016, Hendriks and Vertegaal 2015). In addition, HA-Sumo2 immunosignal was also observed in MAP2-positive regions and, strikingly, in synapses positives for Synapsin1, Shank2, Gephyrin, and/or vGlut1 (Figure 3C and D). Higher magnification images confirmed the presence of Sumo2 along MAP2 dendrites and at synapses (Figure 4; Figure 4–figure supplement 1). Quantification of the 3D object-based co-localization between the anti-HA immunosignal with the various synaptic markers confirmed the presence of Sumo2 at synapses, with HA-Sumo2 signal being significantly co-localized with Shank2, Synapsin 1, vGlut1 and vGAT (Figure 4C and Figure 4–figure supplement 1, which shows a typical 3D reconstruction generated in Imaris). Co-localization of Sumo2 with Gephyrin was not obvious, most likely due to difficulties in immunolabeling inhibitory synapses with the method optimized for HA visualization. Altogether, our data demonstrate the presence of Sumo2 at synapses.

**Figure 3:**
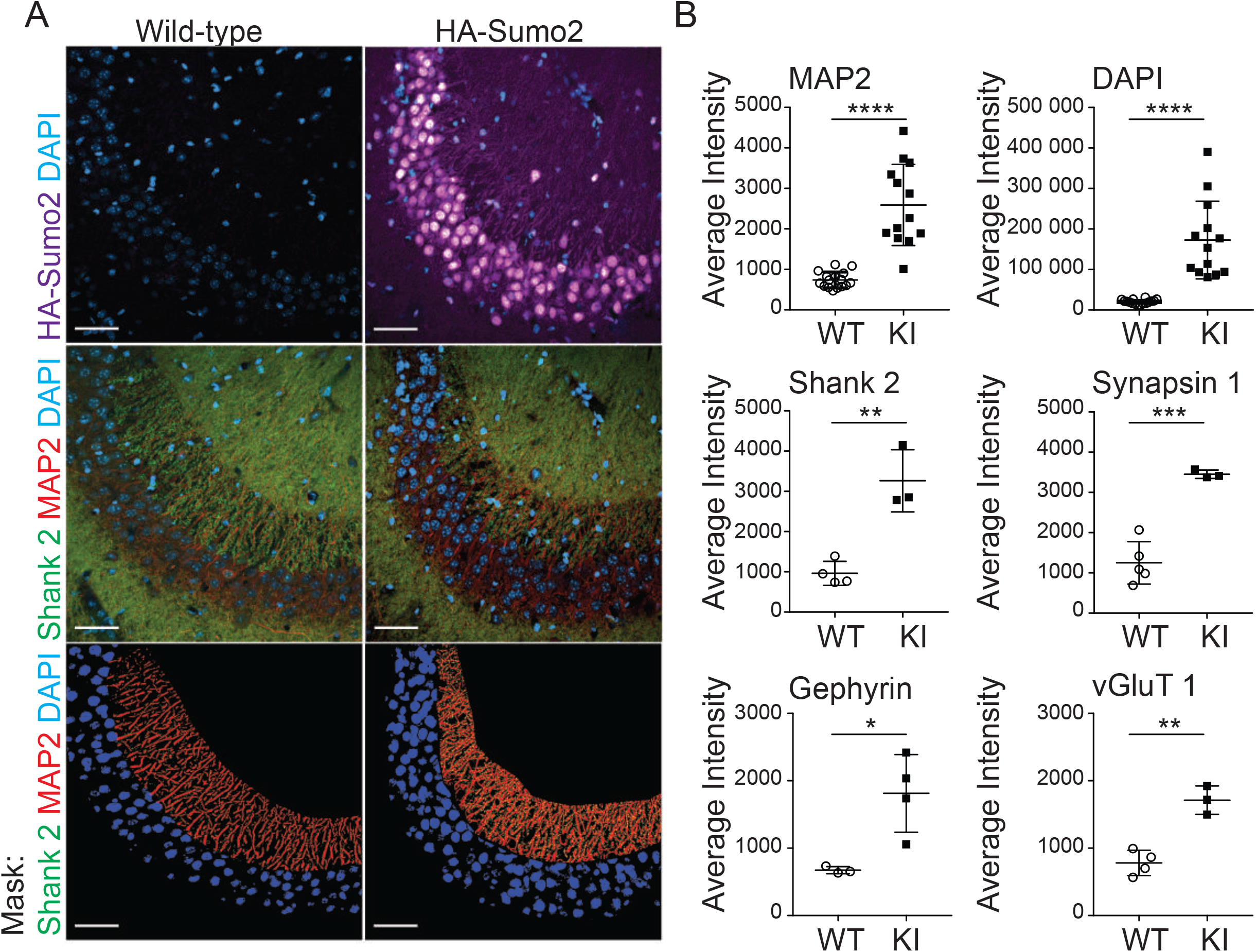
Extra-nuclear localization of Sumo2 in neurons of the hippocampal CA3 region. (A). Anti-HA (purple), DAPI (blue), MAP2 (red), and anti-Shank2 (green) immunostaining of wild-type (left) and HA-Sumo2 KI (right) hippocampal CA3 region. Scale bar: 50 µm. (B) Anti-HA signal in masked regions for each marker (exemplified for Shank2 in the bottom panel A) was quantified in both wild-type (WT) and HA-Sumo2 KI, and average intensity was calculated. N=3, *p<0.05, **p<0.005, ***p<0,0005, ****p<0,0001.

**Figure 4:**
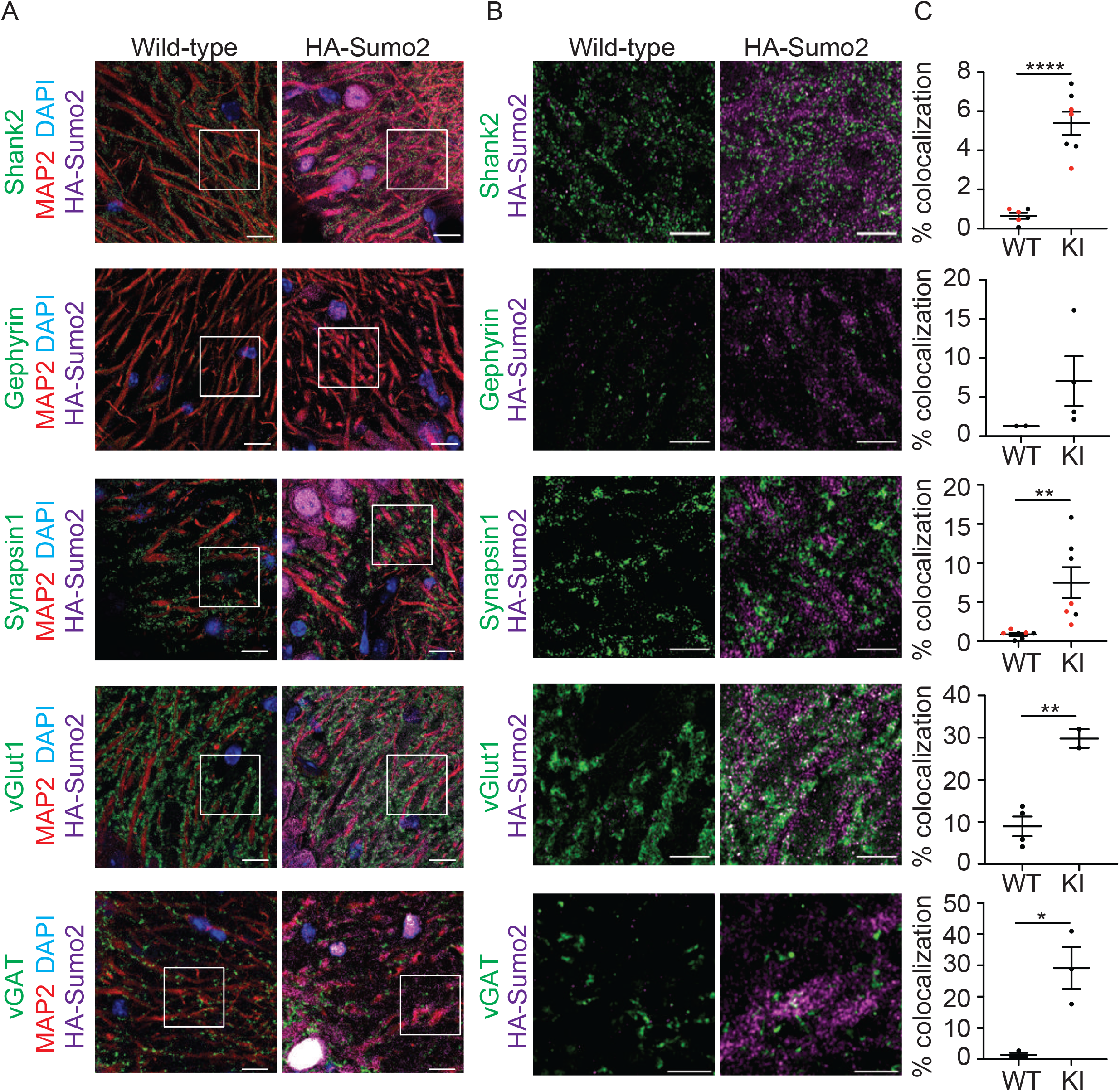
Sumo2 localizes at synapses *in vivo*. (A). High magnification images of wild-type (left) and HA-Sumo2 KI (right) hippocampal CA3 region labelled with anti-HA (purple), DAPI (blue), MAP2 (red), and synaptic markers (green) Shank2, Gephyrin, Synapsin1, vGlut1 and vGAT1. Scale bar, 10 µm. (B). High magnification images with equally scaled anti-HA intensity corresponding to the insets (white boxes) in A showing anti-HA (purple) immunostaining of wild-type (left) and HA-Sumo2 KI (right) hippocampal CA3 region immunolabelled with the synaptic markers (green) Shank2, Gephyrin, Synapsin1, vGLUT1 and vGAT1. Scale bar: 5 µm. N=3. (C). The percentage of co-localization between each synaptic marker and the anti-HA signal was quantified in both wild-type (WT) and HA-Sumo2 KI. N=3, *p<0.05, **p<0.005, ****p<0,0001.

### HA-Sumo mice reveal convergent and contrasting neuronal Sumo1 and Sumo2 substrates

The observation that there are regional and subcellular differences between Sumo1 and Sumo2 indicate divergent roles for the Sumo paralogs in the mouse brain. To uncover such molecular differences between Sumo paralogs, we performed HA-tag immunoprecipitation, under denaturing conditions to break apart standard protein-protein interactions, from whole brain lysate of WT, His_6_-HA-Sumo1, and His_6_-HA-Sumo2 mice, followed by mass spectrometric analysis to identify candidate targets of Sumo1 and Sumo2 *in vivo* (Figure 5A; Table 2). To rank proteins identified from the mass spectrometry dataset, we summated the total peptides from 4 replicates per genotype and then filtered proteins with >2-fold enrichment over proteins identified in the WT condition. Next, to determine high-stringency candidates, proteins were further filtered based on peptide counts being identified in at least 2/4 replicates in the HA-Sumo conditions, and no more than 1/4 replicate in the respective WT condition. Using these criteria, we identified 131 proteins enriched in the HA-Sumo1 and 75 proteins enriched in the HA-Sumo2 immunoprecipitations (Table 2). Gene ontology and Reactome terms related to protein SUMOylation process (Figure 5B) were significantly enriched in both datasets (Figure 5C; Figure 5—figure supplement 1C; Table 3). Indeed, we identified critical components of the SUMOylation pathway including E1 SUMO ligases Sae1 and Uba2, the E2 SUMO ligase Ube2i, and E3 ligases Ranbp2(Pichler et al., 2002), Trim28(Liang et al., 2011), and Pml(Chu and Yang, 2011), validating the use of our mouse model to identify molecular substrates involved with SUMOylation.

**Figure 5:**
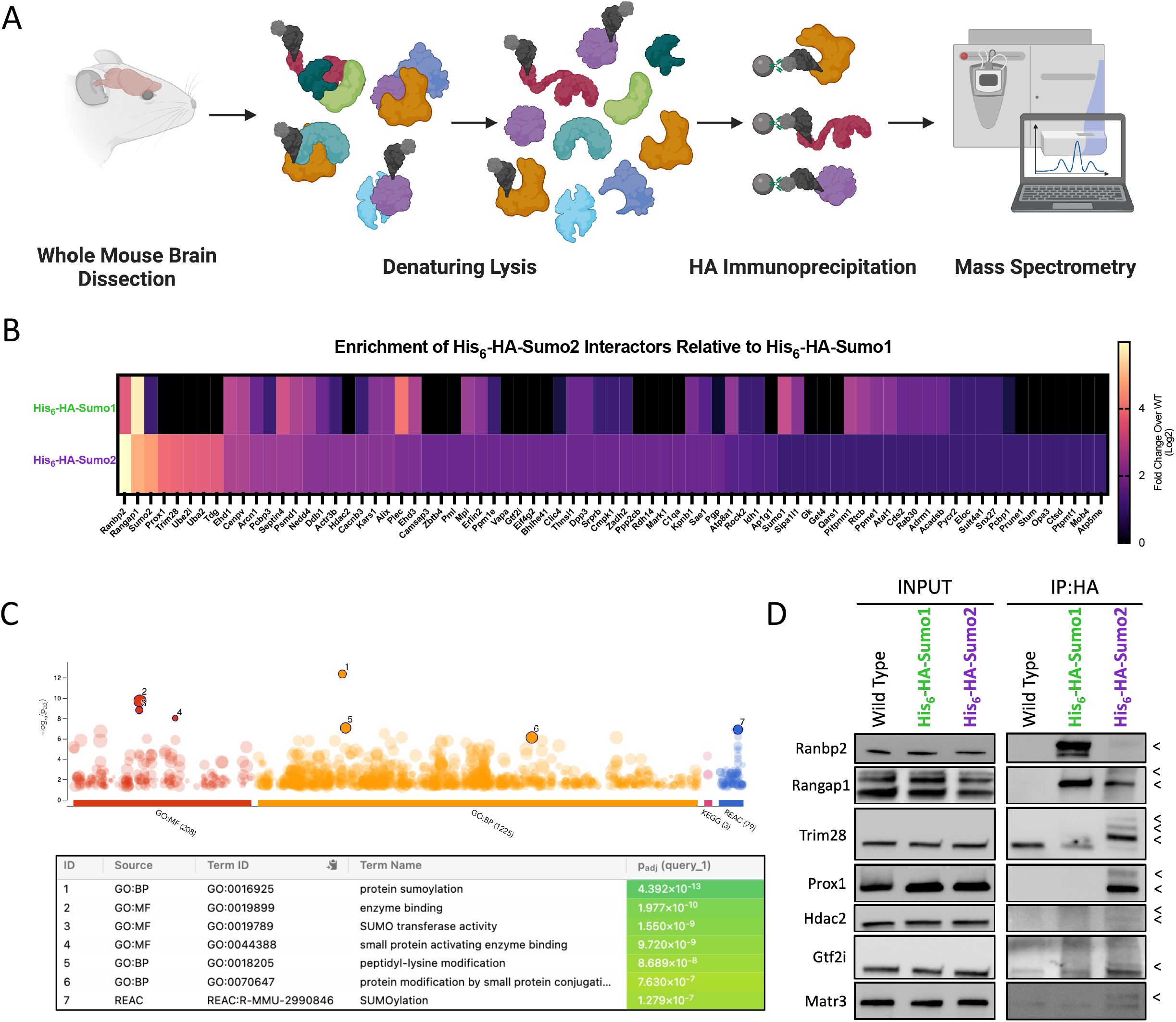
Neuronal Sumo2 has shared and distinct substrates compared to Sumo1 *in vivo*. (A). Schematic of approach to identify His_6_-HA-Sumo interactors from the adult mouse brain via anti-HA affinity immunoprecipitation and mass spectrometry (N=4). (B). Heat map depicting the relative peptide abundance of His_6_-HA-Sumo2 interactors relative to levels in His_6_-HA-Sumo1 immunoprecipitation. (C). gProfiler2 Gene Ontology analysis for His_6_-HA-Sumo2 interactors. (D). Anti-HA affinity Immunoprecipitation followed by Western Blot analysis of His_6_-HA-Sumo1 and His_6_-HA-Sumo2 interactors as listed on the left side (N=3-6). Black arrowheads indicate protein SUMOylation inferred by shift in molecular weight.

To determine whether we could identify specific interactors of Sumo2, we compared the relative fold change of peptides identified in the His_6_-HA-Sumo2 immunoprecipitation to that of the His_6_-HA-Sumo1 immunoprecipitation and found that there are shared, specific, and preferential interactors of Sumo2 vs. Sumo1 (Figure 5D; Figure 5—figure supplement 1D; Table 2). We selected several hits from our mass spectrometry screen for validation via Western blot (Figure 5D). Ranbp2 and Rangap1 are consistently identified amongst the most abundantly SUMOylated proteins (Gareau et al., 2012; Geiss-Friedlander and Melchior, 2007; Saitoh et al., 1998) and were indeed validated to be shared targets of Sumo1 and Sumo2 in the mouse brain. Additionally, both proteins displayed a preference for Sumo1 modification in the mouse brain as previously described (Gareau et al., 2012). Conversely, the E3 ligase and transcriptional repressor Trim28 displayed a preference for Sumo2 modification in the adult mouse brain. Interestingly, the transcription factor Prox1, involved in neurogenesis and a marker of granule cells in the dentate gyrus and cerebellum (Lavado et al., 2010), was one of the most abundantly SUMOylated proteins targeted specifically by Sumo2 in our screen. We further tested the sensitivity of the model to detect more modestly SUMOylated proteins targeted by Sumo2 in the mouse brain, as measured by relative abundance in our immunoprecipitation coupled with mass spectrometry experiment. We confirmed that both Histone Deacetylase 2 (Hdac2) and General Transcription Factor 2i (Gtf2i, also known as Tf2-i) were Sumo2-modified (Figure 5D). Because of the dynamic nature of SUMOylation and the fact that Sumo only targets a fraction of the total pool of most substrates, biologically meaningful interactions may be buried under strong, constitutively SUMOylated targets in our screen. As a proof of concept, we picked a hit with low enrichment in the His_6_-HA-Sumo2 immunoprecipitation by expanding the filtering criteria to a >1.5 fold enrichment over wild-type: Matrin3 (Matr3). We found that Matr3 was selectively SUMOylated in the His_6_-HA-Sumo2 pulldown (Figure 5D), indicating that even at milder cut-off thresholds, SUMOylated substrates identified by mass spectrometry can be validated in an orthogonal assay. Taken together, these results show that the His_6_-HA-Sumo2 mouse model is effective to identify SUMOylated proteins, including proteins SUMOylated at relatively low levels *in vivo*, and can be used for targeted studies of protein substrates.

### Subcellular Localization of Sumo Interaction in Neurons

Sumo1 and Sumo2 predominantly reside in the nucleus, and unsurprisingly, many of the interactors identified here and in other studies of protein SUMOylation are nuclear proteins. However, previous studies hinted at roles for SUMOylation outside of the nucleus(Hasegawa et al., 2014; Ilic et al., 2022; Wang et al., 2014a; Watanabe et al., 2008; Yang and Paschen, 2015). During the validation of whole brain imaging, we observed that His_6_-HA-Sumo2, while extensively localized throughout the nucleus, could also be observed within the somata and synapses of neurons *in vivo*, contrasting the predominantly nuclear membrane staining of His_6_-HA-Sumo1 (Figure 2—figure supplement 2; Figure 3). The cytoplasmic localization of His_6_-HA-Sumo2 in neurons indicates unique extranuclear roles *in vivo*. Indeed, we identified several predominantly extranuclear proteins via mass spectrometry in the anti-HA affinity purification from the His_6_-HA-Sumo2 knock-in mice. To better visualize the subcellular localization of Sumo, we cultured primary cortical neurons from heterozygous His_6_-HA-Sumo1, His_6_-HA-Sumo2, and WT animals and co-stained them for HA and Map2 as a cytoplasmic marker. As previously observed, Sumo2 is predominantly distributed uniformly throughout the nucleus except for heterochromatic foci, but is also observed at low levels in the cytoplasm (Figure 6––figure supplement 1A), contrasting with the anti-HA immunosignal observed for Sumo1, which predominantly localizes at the nuclear membrane. This indicates that Sumo2 specifically may play a larger role in regulating proteins outside of the nucleus.

To better assess the subcellular localization of neuronal SUMOylated proteins targeted by Sumo2, and to provide an additional level of validation in an orthogonal system, we used proximity ligations assays (PLA) against endogenous Sumo2/3 and targets of Sumo2 identified from our mass spectrometry screen to visualize interactions within wild-type primary cortical neurons (Figure 6A) (Ristic et al., 2016; Sahin et al., 2016). In this assay, proteins colocalizing within 40 nm will produce a PLA signal suggesting a potential interaction. We selected proteins expected to interact with Sumo2/3 in the nucleus, Gtf2i and Matr3, as well as expected cytoplasmic interactors Rangap1, Ctsd, Alix, and Kars1 for additional validation (Figure 6B & C). Gtf2i is distributed almost exclusively throughout the nucleus and interactions between Sumo2/3 and Gtf2i, inferred by the presence of PLA foci, are seen throughout the nucleoplasm. Interestingly, PLA foci were often observed to cluster around constitutive heterochromatin (Figure 6C, inset), indicating a potential role for Sumo2 in regulating Gtf2i at the periphery of heterochromatin in neurons. Matr3 is also localized throughout the nucleus and to a lesser extent in the cytoplasm, consistent with its identification in cytoplasmic processing bodies (Rajgor et al., 2016). Interestingly, PLA signals for Matr3 and Sumo2/3 are robust, indicating extensive interactions between the two proteins. Immunofluorescence staining of Rangap1, Ctsd, Alix, and Kars1 demonstrate these proteins predominantly reside outside of the nucleus (Figure 6C). PLA between these hits and Sumo2/3 display an increase in the proportion of cytoplasmic foci relative to nuclear foci (Figure 6B), in addition to a substantial decrease in total nuclear PLA foci compared to nuclear localized proteins Gtf2i and Matr3 (Figure 6—figure supplement 1B). Importantly, for all PLA assays conducted, negative controls did not elicit a PLA signal (Figure 6—figure supplement 1C). Taken together, our PLA analyses of wild-type neurons provide an additional layer of validation of Sumo2-substrate interactions. With the added benefit of providing spatial resolution to the interactions, PLA data support a role for neuronal Sumo2 outside the nucleus.

**Figure 6:**
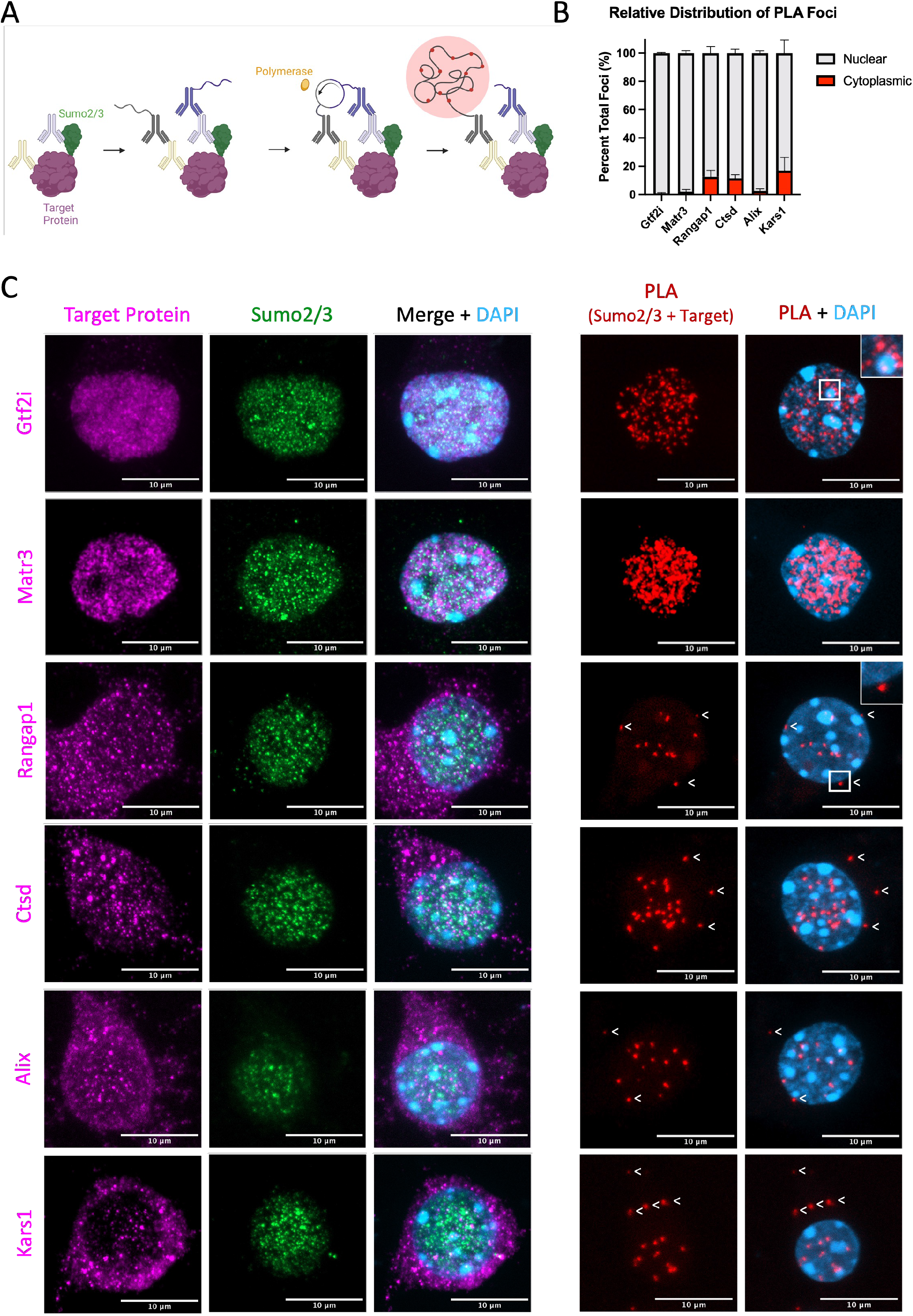
Established and newly identified Sumo2 substrates are present both in nuclear and extranuclear compartments in neurons. (A). Schematic of the proximity ligation assay (PLA) strategy for native Sumo2/3 and target proteins in WT primary cortical neurons. (B). The relative proportion of PLA foci between the selected target proteins and Sumo2/3 was quantified and normalized within the nucleus (grey) versus outside of the nucleus (red) (N=3). (C) Representative Z-projected immunofluorescent images and PLA assays for selected target proteins identified from the mass spectrometry screen (N=3). White arrowheads indicate cytoplasmic PLA foci. Scale bar: 10 μm.

## DISCUSSION

Protein SUMOylation plays a variety of essential and conserved roles (Ilic et al., 2022). Increasing evidence indicates that SUMOylation plays roles in the brain, e.g. in neuronal development or learning and memory(Ripamonti et al., 2020). Furthermore, SUMOylation has been linked with various brain disorders, including Autism Spectrum Disorder, Epilepsy, Schizophrenia, and neurodegenerative diseases such as Alzheimer’s disease, Parkinson’s disease, and Amyotrophic Lateral Sclerosis (Bernstock et al., 2018; Krumova and Weishaupt, 2013; Osmanovic et al., 2022; Rousseaux et al., 2016). However, despite a clear connection with neuronal function and dysfunction, SUMOylation has remained enigmatic due to technical challenges and limitations for studying this modification (Daniel et al., 2017). We generated His_6_-HA-Sumo2 and HA-Sumo2 KI mouse alleles to overcome these technical limitations and to provide insight into the role of native SUMOylation *in vivo*. These mice complement the previously generated His_6_-HA-Sumo1 mouse line (Tirard et al., 2012), which together enable direct comparisons of Sumo1 and Sumo2 *in vivo* to further understand the specific roles conferred by Sumo2 and native roles of SUMOylation.

Using whole brain imaging against the HA epitope in His_6_-HA-Sumo1 and His_6_-HA-Sumo2 mice, we found that Sumo1 and Sumo2 are broadly distributed throughout the mouse brain. Deeper anatomical analysis revealed clear patterns of Sumo differences across brain structures as well as some divergence of Sumo1 and Sumo2 levels in the brain. Indeed, we observed that levels of His_6_-HA-Sumo1 was generally evenly distributed amongst cortical layers, whereas His_6_-HA-Sumo2 levels were higher in layers 2/3 (Figure 2—figure supplement 1D). In the hypothalamus, we found that Sumo levels vary across anatomical regions (Figure 2—figure supplement 1E). Interestingly, regions involved in circadian rhythm including the subparaventricular zone (SBPV) and suprachiasmatic nucleus (SCH) show high levels of Sumo2. These findings are consistent with previous reports that have linked both Sumo1 and Sumo2 differentially affecting circadian clock related proteins such as PER2 and CLOCK, ultimately supporting a role for SUMOylation in regulating circadian rhythm processes (Chen et al., 2021; Drunen and Eckel-Mahan, 2021; Lee et al., 2015).

We observed differential subcellular distribution patterns between Sumo1 and Sumo2. Western blot analysis of brain subcellular fractions showed that Sumo2 is more abundantly expressed than Sumo1 (Figure 1), and that Sumo2 conjugates are present outside of the nuclear compartment, including synaptic fractions (Figure 3–figure supplement 1B & C). This is of importance, as the role of Sumo proteins at synapses has been debated, particularly in the context of Sumo1(Daniel et al., 2018; Daniel et al., 2017). We previously showed that Sumo1 conjugates are not present in synaptic compartments, in contrast to several other reports. Here, we used anti-HA immunolabeling of brain material from HA-Sumo2 mouse brains, and used WT littermates as negative controls to assess non-specific anti-HA immunolabeling (Figures 3, 4 and Figure 4–figure Supplement 1). Strikingly, the quantification of anti-HA immunosignal intensity confirmed a co-localization of Sumo2 with pre- and post-synaptic markers, at both excitatory and inhibitory synapses (Figures 3A-B, 4A-C). This comparative and quantitative approach of investigating the HA-Sumo mouse models allows for the specific assessment of Sumo2 immunosignals within various neuronal sub-compartments, and we provide here the first robust dataset identifying the presence of a Sumo paralog, Sumo2, at synapses in the mammalian brain. These data suggest that Sumo1 and Sumo2 do not only have divergent patterns of expression in mouse brain, but also show differential distributions in the various subcellular compartments, with only Sumo2 present in synapses.

Sumo2 is the only essential Sumo paralog, and has key roles in neurodevelopment, nerve cell function, and neurological diseases(Krumova and Weishaupt, 2013; Ripamonti et al., 2020; Stankova et al., 2018; Wang et al., 2014b). To uncover some of the unique clientele targeted by Sumo2, we performed HA tag immunoprecipitation paired with mass spectrometry. As SUMOylation results in the covalent interaction between Sumo and the target substrate, mouse brains were lysed under denaturing conditions to promote the dissociation of protein complexes prior to immunoprecipitation to help identify bona fide SUMOylation substrates. Using stringent criteria, 75 proteins were identified to potentially interact with Sumo2, ∼1/2 of which were selective to Sumo2. Conversely, using the same criteria, we identified 131 proteins that interact with Sumo1, 98 of which were selective to Sumo1. Pathway analysis of the Sumo2 dataset revealed that gene ontology terms related to SUMOylation were significantly enriched in the dataset, validating this method to identify potentially SUMOylated substrates. Moreover, pathway analysis of the Sumo1 dataset revealed enrichment of proteins involved in protein transport consistent with roles for Sumo1 in nucleocytoplasmic shuttling (Pichler et al., 2002; Salinas et al., 2004; Westman et al., 2010). Prox1 was validated via Western blot as one of the strongest interactors specific to Sumo2 in the mouse brain. Interestingly, previous studies demonstrated that SUMOylation of Prox1 by Sumo1 occurs *in vitro* and in cancer cell lines (Pan et al., 2009; Shan et al., 2008). During development, Prox1 was found to be SUMOylated by Sumo1, and this interaction is critical for neurogenesis in the neural tube development. However, the role of Prox1 in the mature brain remains unclear, although some data indicate an involvement in adult neurogenesis (Correa-Vázquez et al., 2021). Due to the extent of the Prox1 and Sumo2 interaction observed in this study, further analysis may uncover key roles of SUMOylated Prox1 in the adult mouse brain. We further found that the RNA binding protein Matr3 interacts with Sumo2 *in vivo*, indicating that even a mild enrichment over wild-type in our dataset may identify *bona fide* biological interactors. Interestingly, many of the enriched computationally-predicted SUMOylation sites in Matr3 (K588 and K843) reside within a structurally disordered region of the protein that also happens to be enriched for mutations causing Amyotrophic Lateral Sclerosis (ALS) (Johnson et al., 2014). Structurally disordered regions are thought to be critically involved in the formation of protein aggregates in neurodegenerative diseases, and given the key role of SUMO in controlling protein solubility and the recent links between SUMOylation and ALS, SUMOylation of Matr3 may provide key insights into Matr3 biology related to this disease. SUMOylation occurs in a context-specific manner and can regulate protein localization within cells. One of the best characterized examples concerns the regulation of nucleocytoplasmic shuttling across nuclear pore complexes (Pichler et al., 2002; Salinas et al., 2004; Westman et al., 2010). However, SUMOylation can also regulate sub-compartmental localizations, such as protein targeting to the nucleolus or to heterochromatin (Andreev et al., 2022; Mo et al., 2002; Rawat et al., 2021). As Sumo2 was observed throughout the nucleus and outside the nucleus *in vivo* and in cultured neurons, we sought to determine the subcellular localization of Sumo2 interactions with hits identified via IP-MS using a PLA-based approach (Figure 6). Gtf2i was SUMOylated almost exclusively in the nucleus but interestingly appeared to cluster around heterochromatin. Gtf2i can function as both a transcriptional activator and repressor in response to various signals. For example, the specific splice isoforms β and Δ undergo changes in subcellular localization to differentially regulate the immediate early gene *c-Fos* in response to growth factor signaling(Hakre et al., 2006). However, roles for Gtf2i in regulating heterochromatin remain elusive. SUMOylation may provide a dynamic mechanism in response to neuronal activity to regulate both targeted expression at the gene level and global transcriptional changes(Niskanen et al., 2015). Finally, interactions between Matr3 and Sumo2/3 occurred broadly throughout the nucleus. However, interactions between Matr3 and Sumo2 in mouse brain occurred at relatively low levels based on immunoprecipitation assays. This discrepancy observed in the nucleus may be explained via non-covalent interactions with Sumo2 through larger protein complexes as Matr3 typically functions in complex with other DNA and RNA binding proteins (Banani et al., 2016; Keiten-Schmitz et al., 2021, Keiten-Schmitz et al., 2020). Extranuclear proteins Rangap1, Ctsd, Alix, and Kars1, despite having a certain level of PLA signal within the nucleus, displayed an increase in the proportion of cytoplasmic PLA foci relative to nuclear PLA foci implying increased interactions with Sumo2/3 outside the nucleus. Indeed, Rangap1 plays roles in nuclear import, but can also be found to some extent within the nucleus (Cha et al., 2015). PLA foci between Rangap1 and Sumo2/3 were often found near the nuclear membrane, likely at nuclear pore complexes (Figure 6C, inset). Ctsd is a lysosomal protease predominantly localizing to lysosomes, thus interactions between Sumo2/3 and Ctsd may suggest that its SUMOylation may play a role in lysosomal metabolism(Nakanishi, 2003). Previous reports have linked Sumo with the apoptosis pathway(Basu-Shrivastava et al., 2022; Besnault-Mascard et al., 2005; Mojsa et al., 2015); here, we observe potential interactions between Sumo and the apoptotic protein Alix further inferring a role for Sumo in the apoptotic process. Finally extranuclear PLA foci could be observed for the lysyl-tRNA Synthetase Kars1, consistent with reports of Sumo interacting with tRNA-related proteins(Chymkowitch et al., 2017; Chymkowitch and Enserink, 2018; Rohira et al., 2013). Together these results suggest that Sumo2 may play a variety of roles outside the nucleus in various subcellular compartments. While PLA assays provide spatial information of putative protein interactions (i.e. interactions occurring within 40 nm), this does not discount indirect interactions (e.g. protein complexes) (Ristic et al., 2016; Sahin et al., 2016). Thus, differentiating between covalent SUMOylation, interactions with SUMO interacting motifs (SIMs), and protein complexes containing Sumo are not possible using this approach. Extensive localization and biochemical analyses, including arginine to lysine mutagenesis to block covalent SUMOylation, should be performed to properly assess how Sumo confers regulation on a specific target throughout the cell.

In sum, the novel (His_6_-)HA-Sumo2 knock-in mouse lines described here represent powerful tools to facilitate the study of protein SUMOylation via Sumo2 *in vivo*. The HA epitope tag allows for specific yet versatile modes to detect and capture Sumo2 in its native context, without altering its function. Furthermore, these mouse models add to the previously generated His_6_-HA-Sumo1 allele, and thus expand the Sumo knock-in mouse toolkit to begin exploring differential SUMOylation in parallel systems. The versatility of these models extends to all other tissue systems, as Sumo proteins are broadly distributed beyond the brain (Hendriks et al., 2018; Uhlén et al., 2015). Beyond basic biology, these alleles can be used to explore disease processes, e.g. by crossing these mouse lines into disease models, thus enabling the study of the Sumo-linked disease proteome (Stankova et al., 2018). Ultimately, these tools will advance our understanding of the essential biological processes and potential disease targets regulated by SUMOylation.

## MATERIALS AND METHODS

### Mouse Husbandry

His_6_-HA-Sumo1 and His_6_-HA-Sumo2 mice, kept on an FVB/N background, were housed with up to 5 mice per cage on a 12 h light–dark cycle. Mice were fed *ad libitum* and all husbandry was performed by the uOttawa Animal Care and Veterinary Services staff. All animal work for the His_6_-HA-Sumo1 and His_6_-HA-Sumo2 mouse lines were done under the breeding (CMMb-3009 and CMMb-3904) protocols approved under the uOttawa Animal Care Committee.

All experiments regarding the HA-Sumo2 KI mice were performed in accordance with the guidelines for the welfare of experimental animals issued by the State Government of Lower Saxony, Germany (LAVES). Animals were hosted in a pathogen free facility at the Max Planck Institute of experimental Medicine and were maintained in groups in accordance with European Union Directive 63/2010/EU and ETS 123 (individually ventilated cages, specific pathogen-free conditions, 21 ± 1°C, 55% relative humidity, 12 h/12 h light/dark cycle). Mice received food and tap water *ad libitum* and were provided with bedding and nesting material. Cages were changed once a week. Animal health was controlled daily by caretakers and by a veterinarian. Health monitoring (serological analyses; microbiological, parasitological, and pathological examinations) was done quarterly according to FELASA recommendations with either NMRI sentinel mice or animals from the colony. The mouse colony used for experiments did not show signs of pathogens. All experiments were performed during a light cycle.

### Mouse Genotyping

Genomic DNA (gDNA)was isolated from small tail samples collected from mouse embryos (primary cortical neuron experiments), pups (pre-weaning), or adult mice (endpoint). His_6_-HA-Sumo2 mice were genotyped using parallel reactions with primers targeting the *His*_*6*_*-HA* knock-in tag and corresponding genomic loci upstream (Forward: 5′- AGGAAGAGAGCGAGAGAGGAA-3′, Reverse: 5′-CACCACCACTACCCATACGA-3′, 224bp product) and downstream (Forward: 5′-TCGTATGGGTAGTGGTGGTG-3′, Reverse: 5′-AGGAGGAGGGGTGGTTATGT-3′, 276bp product) of the knock-in. 25µL reactions were prepared using <10ng gDNA, primers (400nM final), 1.25µL DMSO (5% final), and 2X GoTaq (Promega). Thermocycler parameters: 95°C for 2 min, (95°C for 30 s, 57.6°C for 30 s, 72°C for 40 s) repeated for 35 cycles, final denature at 72°C for 5 min. HA-Sumo2 mice were genotyped with primers flanking the *HA* knock-in tag (Forward: 5′-GCCCTCTGCCTCGTCCAC-3′, Reverse: 5′-CCGCCGCGAGCTCACCTTG -3′, WT Allele: 160bp product, KI Allele: 187bp product). 20µL reactions were prepared using 1µL clean gDNA, primers (4 pmol final), and 5X Biozym Hot-Start Taq DNA Polymerase plus extra Mg^2+^ (Biozym 331620XL). Thermocycler parameters: 96°C for 3 min, (94°C for 30 s, 62°C for 60 s, 72°C for 60 s) repeated for 32 cycles, final denature at 72°C for 7 min. His_6_-HA-Sumo1 mice were genotyped as previously described (Tirard et al., 2012).

### Biochemical Analysis of Adult Mouse Brain and Spinal Cord

Mice were anesthetized using isoflurane inhalation and sacrificed via decapitation. Brains were quickly isolated, dissected for regional protein analysis, and flash frozen on dry ice. Analysis of total Sumo1 and Sumo2/3 levels were performed on whole brain lysates. Spinal cords were removed using hydraulic extrusion(Richner et al., 2017). Samples were thawed and immediately lysed using a dounce homogenizer in RIPA buffer (9.1 mM dibasic sodium phosphate, 1.7 mM monobasic sodium phosphate, 150 mM sodium chloride, 1 % NP-40, 0.5 % sodium deoxycholate, 0.1 % SDS) containing 50 mM freshly prepared N-Ethylmaleimide (Sigma), 0.25 % β-mercaptoethanol, and Xpert Protease (GenDEPOT) and Xpert Phosphatase (GenDEPOT) inhibitor cocktails. Whole brain lysates were lysed in 11 mL of RIPA buffer described above and centrifuged at 125,000 x g for 2 hours at 4 °C. Regional lysates were centrifuged at 21,000 x *g* for 20 minutes at 4 °C and the supernatant was removed and suspended in 4X laemmli buffer (BioRad) with 2-mercaptaethanol (BioRad) and boiled at 95 °C for 5 minutes. Samples were run on 8 % polyacrylamide gels and transferred onto 0.45µm nitrocellulose membranes at 340 mA for 90 minutes and stained with ponceau to normalize protein levels between lysates. Normalized lysates were run on a 4-15 % GTx Mini-PROTEAN gel (BioRad) and transferred onto a 0.45 µm nitrocellulose membrane at 340 mA for 2 hours for western blot analysis. Densitometry was performed using BioRad Image Lab software and FiJi/ImageJ by normalizing the HA intensity to ponceau for High Molecular Weight (>70 kDa) and free Sumo levels.

### Real Time Quantitative PCR (RT-qPCR) analysis

RNA was extracted from mouse brain homogenate using Trizol-Chloroform extraction (Invitrogen™ User Guide: TRIzol Reagent version B.0). Briefly, mouse brains were homogenized in 3 ml of PEPI Buffer [5 mM EDTA, 1X protease inhibitor (GenDEPOT cat# P3100-020), in 1X PBS] using a dounce homogenizer. 3 % of homogenate was added to 1 ml of TRIzol Reagent (Fisher Scientific cat# 15-596-026) and RNA was isolated as per the user guide referenced above. cDNA was synthesized using 5X All-in-One RT Master Mix (Bio Basic cat# HRT025-10) following manufacturer’s instructions. RT-qPCR was performed using Green-2-Go qPCR Master Mix (Bio Basic cat# QPCR004-S) with 25 ng cDNA per reaction and primers targeting mouse *Sumo1* (NM_009460.2) (Forward: 5′- GCTGATAATCATACTCCGAAAGAAC-3′, Reverse: 5′-CCCCGTTTGTTCCTGATAAA- 3′), *Sumo2*(NM_133354.2) (Forward: 5′-GGACAGGATGGTTCTGTGGTGC-3′, Reverse: 5′-CCCATCAAACCGGAATCTGATCTGC-3′), and *Hprt1*(NM_013556.2) (Forward: 5′- TGATAGATCCATTCCTATGACTGTAGA-3′, Reverse: 5′- AAGACATTCTTTCCAGTTAAAGTTGAG-3′). Reactions were run on BioRad CFX96 thermocycler (protocol: 95°C for 5 min, 40 cycles of 95°C for 15 s and 60°C for 60 s, then melting curve). *Sumo1* and *Sumo2* Ct values were standardized to the Ct values of *Hprt1*.

### Brain preparation for whole brain clearing and HA immunolabeling

Mice were sedated via intraperitoneal injection of 16.25 mg sodium pentobarbital (Euthanyl, DIN 00141704). Once sedated, mice were perfused using 10 mL 1X PBS + 10 U/mL heparin (Millipore Sigma, H3393-50KU) followed by 10 mL of freshly prepared 4 % paraformaldehyde. Brains were carefully isolated and stored in 13 mL of 4 % paraformaldehyde at 4 °C overnight with gentle shaking. Brains were rinsed with 1X PBS then shipped to LifeCanvas Technologies (MA) in 1X PBS + 0.02 % sodium azide.

### Whole mouse brain processing, staining, and imaging

Whole mouse brains were processed using the SHIELD protocol (LifeCanvas Technologies (Park et al., 2018)). Samples were cleared for 1 day at 42 °C then actively immunolabeled using SmartBatch+ (LifeCanvas Technologies) based on eFLASH technology integrating stochastic electrotransport (Kim et al., 2015) and SWITCH (Murray et al., 2015). Each sample was labeled with 60 µg anti-NeuN (Encor, MCA-1B7) and 36 µg rabbit anti-HA-tag (Cell Signalling Technology #3724) followed by fluorescently conjugated secondary antibodies in a 3:2 primary:secondary molar ratio (Jackson ImmunoResearch). Samples were incubated in EasyIndex (LifeCanvas Technologies) for refractive index of RI = 1.52 and imaged at 3.6X using a SmartSPIM axially-swept light sheet microscope (LifeCanvas Technologies). Images were tile corrected, de-striped, and registered to the Allen Brain Atlas (Allen Institute: https://portal.brain-map.org/). NeuN channels for each brain were registered to 8-20 atlas-aligned reference samples using successive rigid, affine, and b-spline warping (SimpleElastix: https://simpleelastix.github.io/). Average alignment to the atlas was generated across all intermediate reference sample alignments to serve as the final atlas alignment value per sample. Fluorescent measurements from the acquired images were projected onto the Allen Brain Atlas to quantify the total fluorescence intensity per region defined by the Allen Brain Atlas. These values were then divided by the volume of the corresponding regional volume to calculate the intensity per voxel measurements.

### HA Immunoprecipitation from mouse brain lysates

Immunoprecipitation was performed based on the previously established protocol(Tirard et al., 2012; Tirard and Brose, 2016). Briefly, mice aged 9-16 weeks old were anesthetized and sacrificed via decapitation and the brain was quickly removed and homogenized using a dounce homogenizer in RIPA buffer (9.1 mM dibasic sodium phosphate, 1.7 mM monobasic sodium phosphate, 150 mM sodium chloride, 1 % NP-40, 0.5 % sodium deoxycholate, 0.1 % SDS) containing 50 mM freshly prepared N-Ethylmalamide (Sigma), 0.25 % β-mercaptoethanol, and Xpert Protease (GenDEPOT) and Xpert Phosphatase (GenDEPOT) inhibitor cocktails. Samples were lysed on ice for 20 minutes with vigorous vortexing every 5 minutes before ultracentrifugation at an average of 100,000 x *g* for 30 minutes at 4 °C. Supernatants were removed and spiked with 50 mM N-ethylmalamide before being added to 50 μL of magnetic protein G Dynabeads (Invitrogen) pre-conjugated with 10 μg mouse HA antibody (in house) per sample. Samples were then placed on rotator at 4 °C for 1 hour. Beads were washed three times in 10 mL of RIPA buffer containing 20 mM N-Ethylmalamide, protease, and phosphatase inhibitors. Wash buffer was thoroughly removed, and beads were eluted using 75 µg synthetic HA peptide (Sino Biological Inc.).

### Sample Preparation for Mass Spectrometry

Two thirds of eluted protein sample from HA Immunoprecipitation were run on a 4-15% Mini-PROTEAN (Bio-Rad) gel to separate proteins and remove synthetic HA peptide. Samples were stained using Silver Stain for Mass Spectrometry Kit (Thermo Scientific, 24600). Lanes were cut into 6-7 gel slices per sample and stored in 1% acetic acid until analysis via LC-MS/MS.

### Protein Identification by LC-MS/MS

Proteomics analysis was performed at the Ottawa Hospital Research Institute Proteomics Core Facility (Ottawa, Canada). Proteins were digested in-gel using trypsin (Promega) according to the method of Shevchenko (Shevchenko et al., 2006), but without the use of iodoacetamide for cysteine alkylation due to the treatment of 50 mM N-ethylmaleimide (Sigma) during the sample immunoprecipitation. Peptide extracts were concentrated by Vacufuge (Eppendorf). LC-MS/MS was performed using a Dionex Ultimate 3000 RLSC nano HPLC (Thermo Scientific) and Orbitrap Fusion Lumos mass spectrometer (Thermo Scientific). MASCOT software version 2.7.0 (Matrix Science, UK) was used to infer peptide and protein identities from the mass spectra. The observed spectra were matched against *Mus musculus* sequences from SwissProt (version 2021-02) and against an in-house database of common contaminants. The results were exported to Scaffold (Proteome Software, USA) for further validation and viewing and will be uploaded to the ProteomXchange Consortium.

### Gene Ontology analysis

Top His_6_-HA-Sumo1 and His_6_-HA-Sumo2 interactors were analyzed using gProfiler2 web tool (https://biit.cs.ut.ee/gprofiler/gost)(Peterson et al., 2020). Lists were arranged in descending order based on relative peptide abundance and analyzed as an ordered query for *Mus musculus* proteins using Benjamini-Hochberg FDR significance threshold with an alpha of 0.05.

### Immunohistochemistry of Mouse Brain

Mice were anesthetized with 30 µl of 120 mg/kg Euthanyl (DIN00141704) and then perfused with 10 ml 1x phosphate buffered saline (PBS) and 10 ml 4 % paraformaldehyde (PFA). Brain and spinal cord tissue were collected and stored for 48 hours in 4 % PFA. Brain and spinal cord tissue were then dehydrated in 10 %, 20 % and 30 % sucrose solutions for 48 hours each before being flash frozen in -40 °C isopentane for 1 minute. Tissues were then sectioned at 20 µm and -21 °C on the Thermo Scientific HM 525 NX cryostat at the Louise Pelletier Histology Core at the University of Ottawa and stored free floating in 1x PBS + 0.02 % NaN3 at 4 °C until use in staining. Brains tissue were incubated for 24 hours in blocking buffer (1.5 % Triton X-100, 5 % cosmic calf serum in 1X PBS), 24 hours in primary antibody (1:500 HA-Tag C29F4 rabbit monoclonal antibody (Cat: 3724S, Cell Signaling Technology), and 1 hour in secondary antibody (1:500 Alexa Fluor 568 donkey anti mouse antibody, Cat: A10037, Lot: 1917938)) with DAPI (1:1,000 Millipore Sigma, D9542-1MG). Tissue was washed 5 times for 5 minutes each in 1x PBS between each treatment and mounted on Fisherbrand Superfrost Plus slides. Slides were left to dry for 24 hours and covered with DAKO mounting medium (Cat: S3023, Lot: 11347938) and #1.5 coverslips.

For the analysis of the synaptic localization of Sumo2, HA-Sumo2 KI brains were used (Figure 1—figure supplement 1B). Mice were anaesthetized (250 mg/Kg Avertin i.p.) and transcardially perfused with PBS and then with 4 % (w/v) paraformaldehyde (PFA) in 0.1 M phosphate buffer (PB), pH 7.4 at 4 °C for 10 min. Brains were removed and post-fixed for 1 h at 4 °C. The tissue was cryoprotected in 30 % (w/v) sucrose in phosphate-buffered saline (PBS). Brains were frozen in the cryostat and sagittal 35 µm sections were prepared with a cryostat and collected free-floating in PBS. For immunohistochemistry, sections were pre-incubated for 24 h in PB containing 3 % horse serum (HS), 3 % fish skin gelatin (FSG), and 0.3 % Triton X-100, and were then incubated for 3 days at 4 °C in primary antibodies diluted in PBS containing 3 % HS, 3 % FSG and 0.3 % Triton X-100. After washing repeatedly in PBS and overnight, sections were incubated overnight in dye coupled secondary antibodies, repeatedly washed, and mounted on slides with Aquapolymount (Polysciences). The antibodies used are listed in Table 4.

### Primary Cortical Neuron Cultures

Pregnant mice were euthanized between at gestation E14.5-15.5 with 48 mg Pentobarbital Sodium (Bimeda-MTC, 8015E) delivered via intraperitoneal injection. Embryos were removed, washed in chilled PBS (Wisent Bioproducts, 311-010-CL), and cortices were isolated in chilled HBSS (Sigma Aldrich, H9394). Cortices were dissociated for 20 minutes with trypsin (Thermo Scientific, 90305) at 37 °C before adding trypsin inhibitor and DNase solution to quench reaction. Cells were pelleted at 2,500 x *g* for 5 minutes at 4 °C and washed with trypsin inhibitor with DNase solution. Cortical neurons were pelleted at 2,500 x *g* for 5 minutes at 4 °C and resuspended in 1 mL Neurobasal outgrowth media (Thermo Scientific, 21103049), supplemented with B-27 (Thermo Scientific, 17504044), N-2 (Thermo Scientific, 21103049), 500 μM L-Glutamine (Wisent Bioproducts, 609-065-EL), and 0.5 % penicillin/streptomycin (GE Healthcare Life Sciences, SV30010) before plating. Cultures were maintained for 7 days in vitro with a half media change after 3-4 days.

### Immunofluorescence in Primary Cortical Neurons

Micro Coverglass #1.5 coverslips (Electron Microscopy Sciences) coverslips were pre-coated with poly-D-lysine (50 μg/ml) overnight at 37 °C, then washed with distilled water three times and air-dried at room temperature for at least two hours. Primary mouse cortical neurons were seeded at 75,000 cells per coverslip were seeded and cultured as described in Primary Cortical Neuron Cultures. On day 7, neurons were fixed using 10 % phosphate buffered formalin for (Fisher Chemical, SF100-4) for 10 minutes followed by 3 × 5-minute washes in 1 mL of 1X PBS. Neurons were blocked in 500 µL of blocking buffer (1.5 % Triton X-100, 10 % cosmic calf serum in 1X PBS) for 1 hour, then incubated in 300 µL of primary antibody diluted in blocking buffer overnight at 4 °C. The following day, the neurons were washed for 4 × 5-minute washes in 1 mL of 1X PBS then incubated in 300 µL of secondary antibody diluted in blocking buffer for 2 hours at room temperature. Next, the neurons were washed for 4 × 5-minute washes in 1 mL of 1X PBS, dried, and then placed on slides with Vectashield Antifade Mounting Medium with DAPI (MJS Biolynx Inc., H-1200). Z-Stack images were obtained on a Zeiss AxioObserverZ1 LSM800 Confocal Microscope at 63X magnification through a Z distance of 10 μm per image using optimal 0.27 μm spacing per slice. The dimensions were set to 1,024 × 1,024 pixels.

### Proximity Ligation Assay in Primary Cortical Neurons

Primary mouse cortical neurons were cultured and fixed as described for immunofluorescence experiments. Individual coverslips for Proximity Ligation Assay experiments (PLA) were transferred to 12-well plates and outlined with a hydrophobic pen. Neurons were blocked using 40 µL of Duolink blocking buffer (Sigma Aldrich, DUO82007) at 37 °C for 1 hour and were then washed for 3 × 5 minutes washes in 1 mL 1X PBS. Next, the neurons were incubated in 300 µL of primary antibody diluted in blocking buffer (1.5 % Triton X-100, 10 % cosmic calf serum in 1X PBS) overnight at 4 °C. The following day, the neurons were washed for 3 × 5 minutes washes in 1 mL of Duolink Wash Buffer A (0.01 M Tris-Base, 0.15 M NaCl, 0.05 % Tween-20, pH 7.4) followed by incubation in 40 µL of Duolink PLA MINUS (Sigma Aldrich, DUO82004) and PLUS probes (Sigma Aldrich, DUO82002) at 37 °C for 1 hour. Duolink PLA probes were diluted in antibody diluent (Sigma Aldrich, DUO82008) at a 1:5 dilution. Neurons were washed for 3 × 5 minutes washes in Duolink Wash Buffer A and then incubated in 40 µL of ligase (Sigma Aldrich, DUO82027) at 37 °C for 30 minutes. Ligase was diluted in 1X ligation buffer (Sigma Aldrich, DUO82009) at a 1:40 dilution. Neurons were washed for 3 × 5-minute washes in Duolink Wash Buffer A and then incubated in 40 µL of polymerase (Sigma Aldrich, DUO82028) at 37 °C for 90 minutes. Polymerase was diluted in 1X amplification buffer (Sigma Aldrich, DUO82011) at a 1:80 dilution. Then, neurons were washed 2 × 10 minutes in Duolink Wash Buffer B (0.2 M Tris-Base, 0.1 M NaCl, pH 7.5) and then again in 1 mL of Duolink Wash Buffer B diluted at 1:100 for 1 minute. Coverslips were briefly air dried and then mounted on slides using Vectashield Antifade Mounting Medium with DAPI. Z-Stack images were obtained on a Zeiss AxioObserverZ1 LSM800 Confocal Microscope at 63X magnification with a 5X digital zoom through a Z distance of 10 μm per image using optimal 0.27 μm spacing per slice with dimensions set to 512 × 512 pixels. Images were analyzed and quantified using the Spots function on the Imaris (ver. 9.9.1 Bitplane, Switzerland) software. Localization of the foci (nuclear versus cytoplasmic) was determined using the Orthogonal Views function for colocalization with DAPI signal.

### Tissue Lysis

Dissected mouse organs were lysed in 1X RIPA buffer (9.1 mM Na_2_HPO_4_, 1.7 mM NaH_2_PO_4_, 150 mM NaCl, 1 % NP-40, 0.5 % sodium deoxycholate, 0.1 % SDS, 1X protease inhibitor (GenDEPOT, P3100), 1X phosphatase inhibitor (GenDEPOT, P3200) and 50 mM N-ethylmaleimide (Sigma-Aldrich, E3876-5G) using a Dounce homogenizer. Tissue lysates were centrifuged at ∼21,000 x *g* for 15 minutes at 4°C and the resulting supernatant was transferred into a fresh microcentrifuge tube.

### Subcellular fractionation

Western blot analysis of brain subcellular fraction was performed as described previously(Jones and Matus, 1974; Tirard et al., 2012).

### Western blot analysis

SDS-PAGE was performed with standard discontinuous gels or with commercially available 4%-12% Bis-Tris gradient gels (Invitrogen). Western blots were probed using primary and secondary antibodies as indicated in Table 4. Blots were routinely developed using enhanced chemiluminescence (GE Healthcare) and imaged using an INTAS ECL Chemostar PLUS Imager HR 6.0. For quantitative Western blotting, transferred proteins were visualized using the total protein stain MemCode (Invitrogen). Western blot signals were visualized using an INTAS ECL Chemostar PLUS Imager HR 6.0 apparatus and quantified using ImageJ, and a ratio of the antibody signal relative to the total protein stain as revealed by the MemCode was performed.

### Confocal Imaging and Image analysis

MAP2, HA-Sumo2, DAPI and the respective synaptic marker fluorescence intensities were acquired from the hippocampal CA3 pyramidal layer and the *stratum radiatum*. In particular, multi-channel z-stacks of 354 × 354 µm-sized fields of view were acquired with a 40x oil immersion objective (1.4 NA) of a Nikon Ti, Yokogawa W1 spinning disk microscope with Andor iXon Ultra 888 EMCCD camera. A custom macro written within the FIJI software package (Schindelin et al., 2012) automatically identified the z-plane of highest HA-Sumo2 intensity, out of which all channels were extracted. The DAPI, MAP2 and synaptic marker channel of this z-plane were segmented with the Trainable WEKA Segmentation FIJI plugin (Arganda-Carreras et al., 2017), resulting in binary masks (Figure 3A, bottom row) that were subsequently used to extract the respective HA-Sumo2 mean intensity.

To more closely investigate HA-Sumo2 colocalization with the respective synaptic markers, high-resolution, multi-channel fields of view (x, y: 30, 30 µm) of the respective stains within the Mossy fiber region were acquired using 63x oil-immersed objective (NA=1.4) on a Leica SP8 confocal laser scanning microscope. Oversampled (x, y, z: 30, 30, 100 nm), 1.2 µm-sized z-stacks within the mossy fiber region were acquired for each fluorescent channel. Signal to noise ratio and resolution were subsequently enhanced and sample geometry was corrected with the SVI Huygens deconvolution software package (ver. 22.04, Hilversum, Netherlands). To obtain object-based colocalization information, an Imaris (ver. 9.9.1 Bitplane, Switzerland) batch process was created to automatically process all three-dimensional fields-of-view identically. All HA-Sumo2 and the respective synaptic marker objects were created, yielding the total object number for the respective stain. Synaptic marker objects were designated as collocating with the HA-Sumo2 objects only if they overlapped with HA-Sumo2 objects and if they contained a minimum HA-Sumo2 average intensity. Data resulting from both analysis workflows was assembled and quantified with the KNIME software package (ver. 4.5.2, Zurich, Switzerland).

### Statistical Analysis

Statistical tests were performed using PRISM 9.4. Test type was picked based on the number of comparisons made. Levels of statistical significance are indicated in figure legends.

## Figure Legends

**Figure 1—figure supplement 1:**
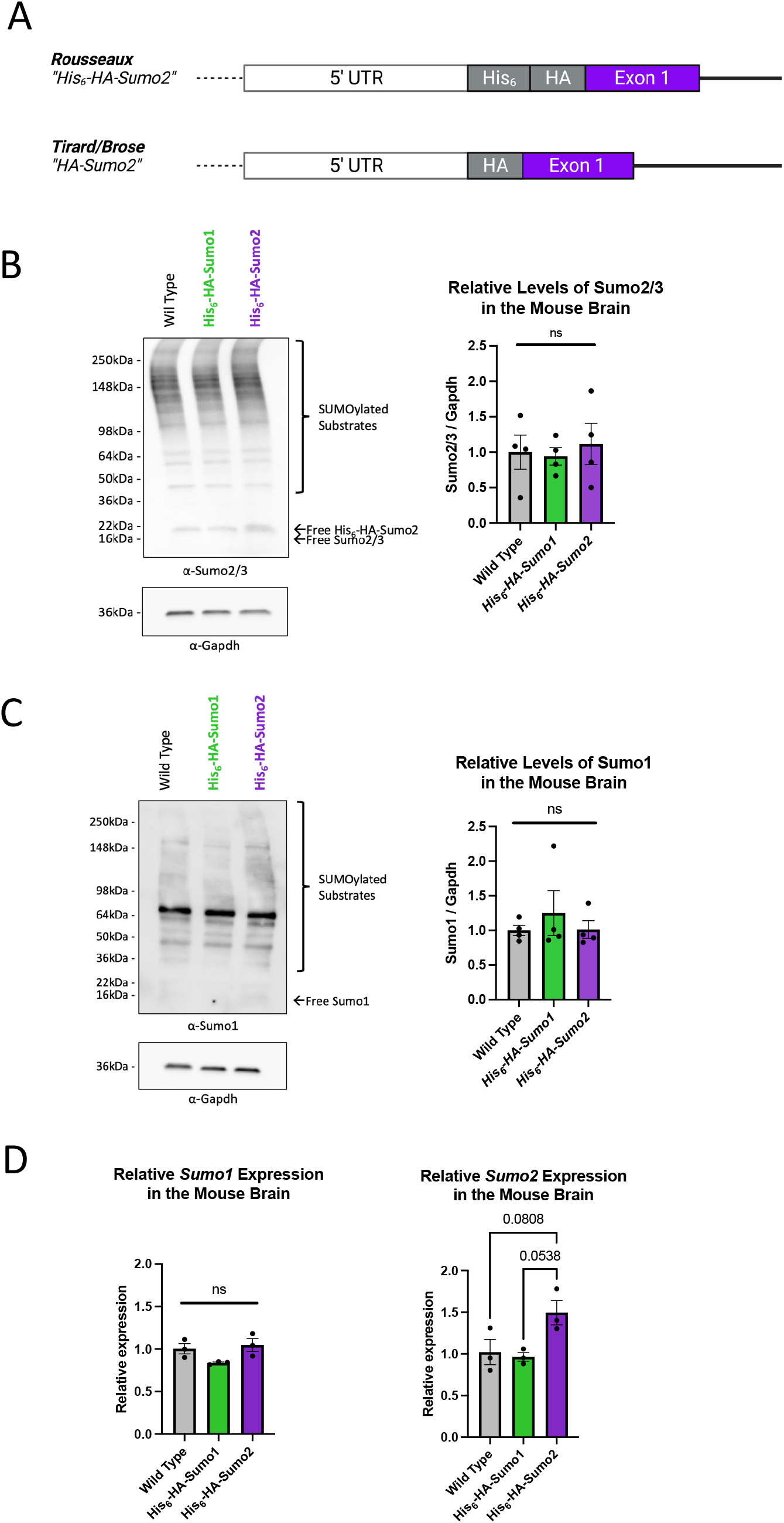
Heterozygous His_6_-HA-Sumo2 and His_6_-HA-Sumo1 mice exhibit normal Sumo levels. (A). Schematic of His_6_-HA-Sumo2 and HA-Sumo2 knock in mice. (B). Anti-Sumo2/3 Western blot analysis of total brain homogenates from heterozygous His_6_-HA-Sumo1, His_6_-HA-Sumo2 mice and WT controls. Anti-Sumo2/3 signal (bracket on the right side of the top left panel) was analyzed by densitometry. Signal from either heterozygous His_6_-HA-Sumo1 or His_6_-HA-Sumo2 mice was normalized to the Gapdh signal and is relative to WT controls (right panel), N=4. (C). Anti-Sumo1 Western blot analysis of total brain homogenates from heterozygous His_6_-HA-Sumo1, His_6_-HA-Sumo2 mice and WT controls. Anti-Sumo1 signal (bracket on the right side of the top left panel) was analyzed by densitometry. Signal from heterozygous His_6_-HA-Sumo1 and His_6_-HA-Sumo2 mice was normalized to the Gapdh signal (bottom left panel) and is relative to WT controls (right panel), N=4. (D). RT-qPCR analysis of *Sumo1* and *Sumo2* transcript levels in whole brain from heterozygous His_6_-HA-Sumo1, His_6_-HA-Sumo2 mice and WT controls. *Sumo1* and *Sumo2* transcript levels were normalized to *Hprt1* as a housekeeping control. All statistical tests were analyzed using an ordinary One Way ANOVA with Tukeys multiple comparisons tests.

**Figure 2—figure supplement 1:**
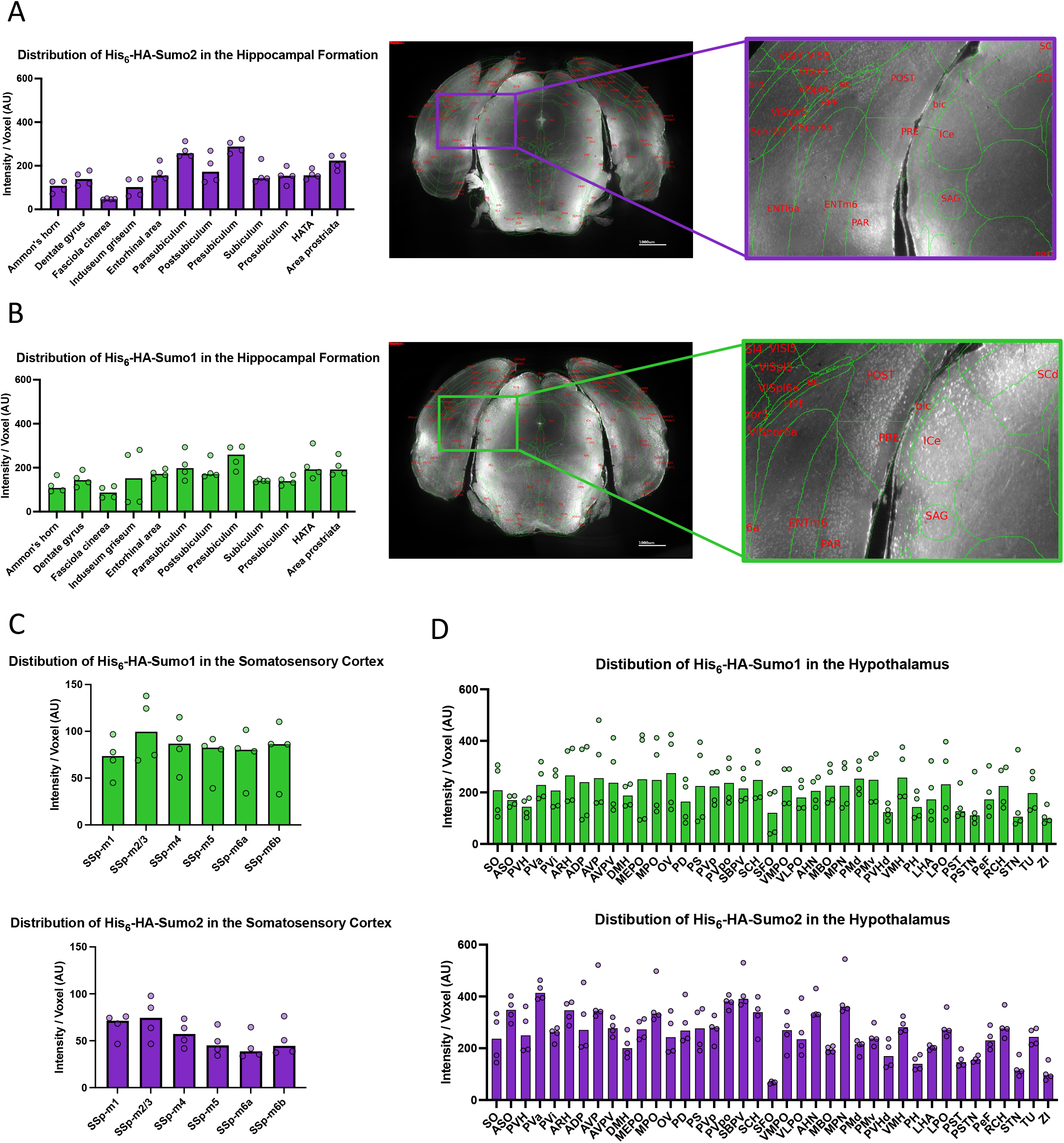
Detailed examination of regional anatomical Sumo paralog distribution. (A). Bar plot depicting the relative levels of anti-HA immunosignal in the regions of His_6_-HA-Sumo2 hippocampus. Each datapoint is the mean intensity from a single hemisphere (N=2, two hemispheres). Middle and Left panels depict a representative image and zoom of the hippocampal formation overlayed with a mask depicting anatomical regions defined by the Allen Brain Atlas. Scale Bar: 1000 µm. (B). Bar plot depicting the relative levels of anti-HA immunosignal across His_6_-HA-Sumo1brain regions. Middle and Left panels depict a representative image and zoom of the hippocampal formation overlayed with a mask depicting anatomical regions defined by the Allen Brain Atlas. Scale Bar: 1000 µm. (C). Bar plot depicting relative anti-HA immunosignal levels in the primary somatosensory area–mouth (SSp-m) layers of His_6_-HA-Sumo1 (Top) and His_6_-HA-Sumo2 (Bottom) brain (D). Bar plot depicting relative anti-HA immunosignal levels in regions of the hypothalamus of His_6_-HA-Sumo1 (Top) and His_6_-HA-Sumo2 (Bottom). Each datapoint from the brain atlases depict the mean intensity from a single hemisphere (N=2, two hemispheres). For each abbreviated brain region, refer to Table 1 for legend.

**Figure 2—figure supplement 2:**
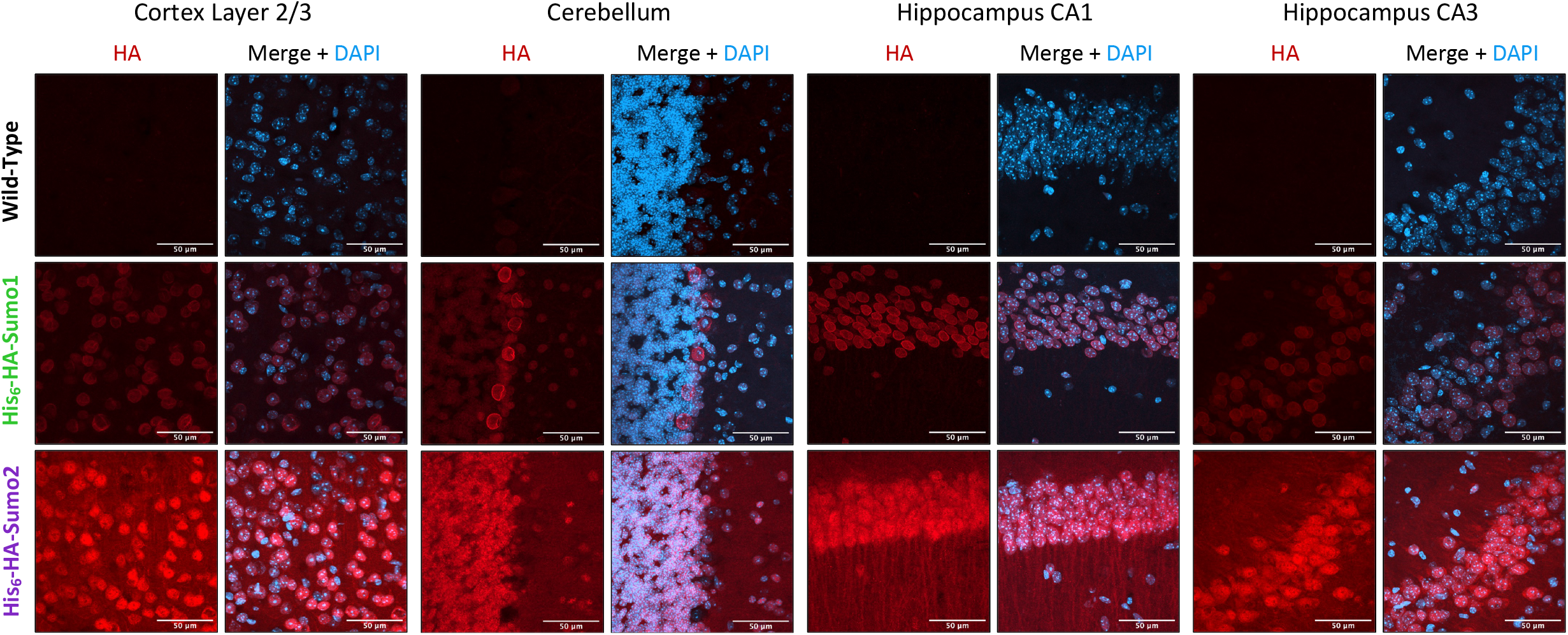
Sumo1 and Sumo2 share similar and distinct anatomical locations and subcellular compartments. Confocal microscopy analysis of anti-HA (red) and DAPI (blue) nuclear labeling of wild-type (untagged, top lane), His_6_-HA-Sumo1 (middle lane) and His_6_-HA-Sumo2 KI (bottom lane) cortex layer 2/3, cerebellum, and hippocampal CA1 and CA3 regions. Z-projection images are representative of three independent replicates. Scale bar: 50 µm

Figure 2—figure supplement video 1: Imaris 3D render of anti-HA immunosignal in whole His_6_-HA-Sumo1 brain. Scale bar: 3000 μm

Figure 2—figure supplement video 2: Imaris 3D render of anti-HA immunosignal in whole His_6_-HA-Sumo2 brain. Scale bar: 3000 μm

**Figure 3–figure supplement 1:**
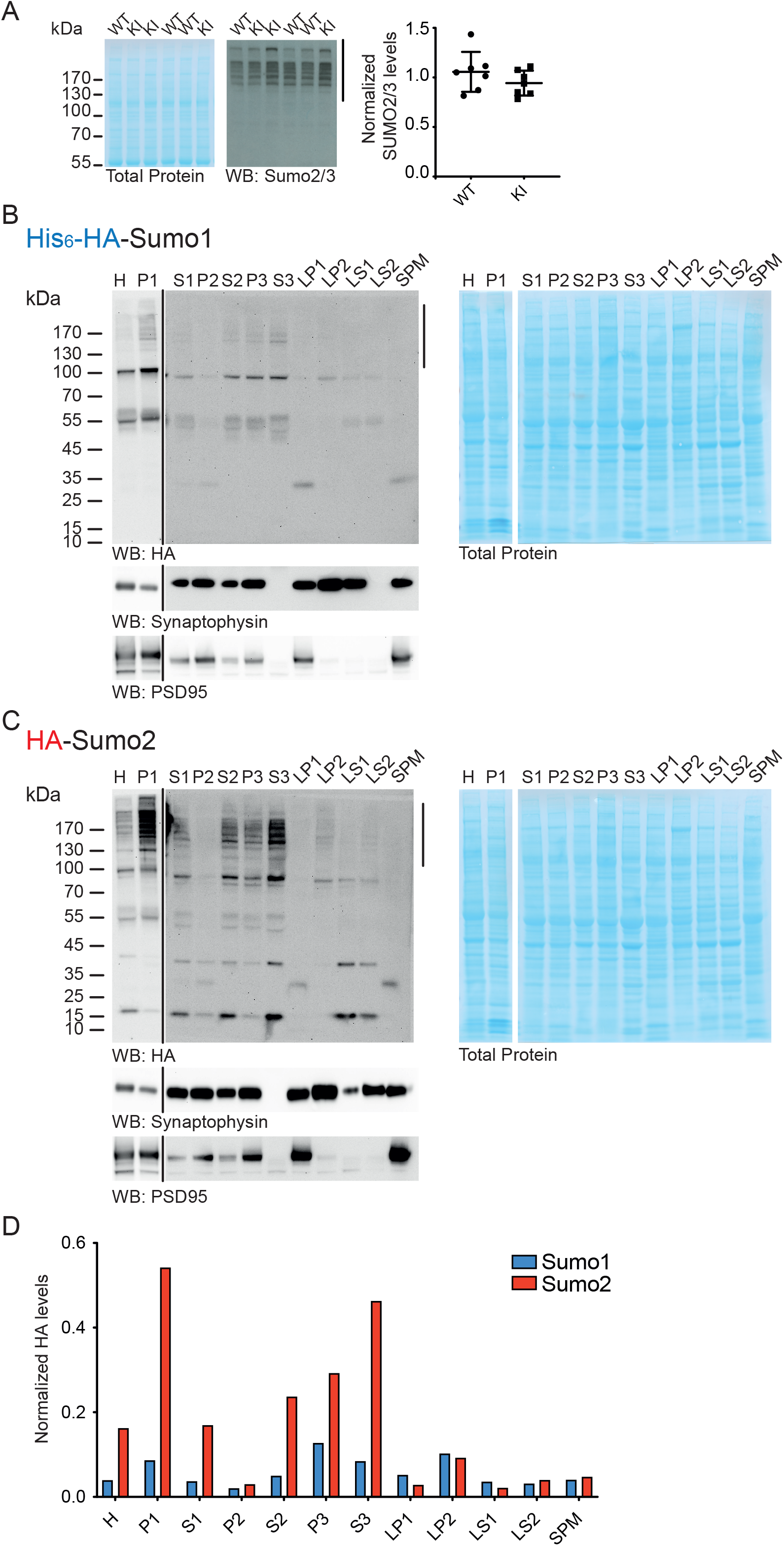
Differential levels and sub-cellular distribution of Sumo1 and Sumo2 conjugates in whole mouse brain fractions. (A). Total Protein stain (left panel) and anti-HA Western blot (right panel) analysis of whole brain lysates from homozygous HA-Sumo2 and corresponding WT littermates. The black bar on the right indicates Sumo2 signal used for the quantification shown by the dot plot (right panel). (B and C). Total protein stain (right panel) and anti-HA (left panel) Western blot analysis of subcellular fractions from homozygous His_6_-HA-Sumo1 (B) and HA-Sumo2 KI (C). Western blot analysis using anti-Synaptophysin and PSD95 (bottom panels) validates the subcellular fractionation procedure. The black line between the HA panel indicates that different exposure times are depicted, as the HA signal in the P1 fraction saturates faster than in the other fractions. The black line on the right of the HA panels indicate the Sumo signal used for quantification in E. H, homogenate; P1, nuclear pellet S1; supernatant after P1 sedimentation; P2, crude synaptosomal pellet; S2, supernatant after P2 sedimentantion; P3, pellet after ultracentrifugation of the S2, cytosolic pellet; S3, supernatant after P3 sedimentation; LP1, lysed synaptosomal membranes; LS1, supernatant after LP1 sedimentation; LP2, pellet after sedimentation of the LS1; LS2, supernatant after LP2 sedimentation; SPM, synaptic plasma membranes. (D). Bar plot depicting the quantification of anti-HA signal as indicated by the black line on the right side of each anti-HA Western blot in C and D, relative to the total protein stain.

**Figure 4–figure supplement 1:**
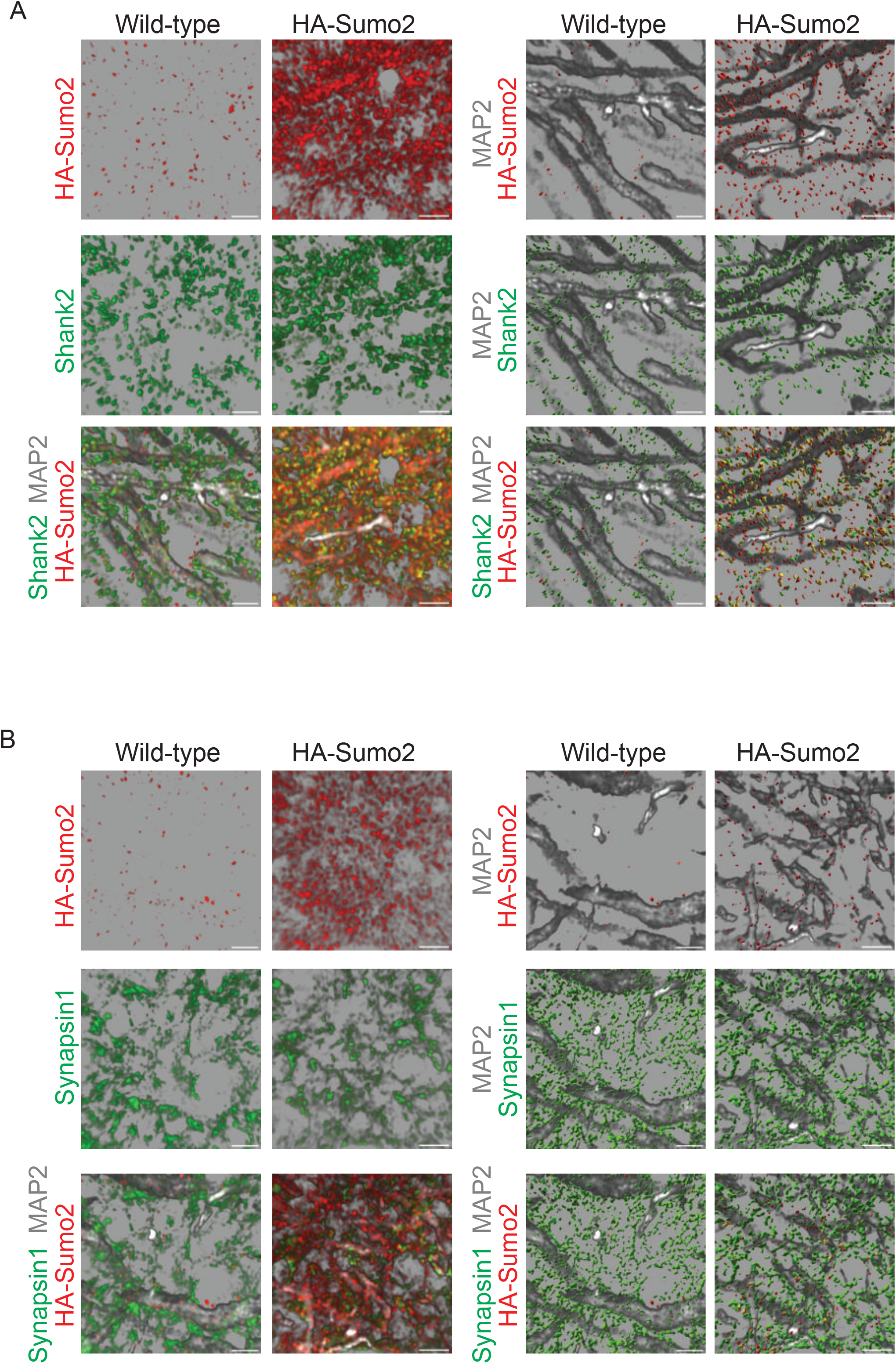
Imaris-based representation of the synaptic Sumo2 co-localization with Shank2 and Synapsin1. Blended representation generated in Imaris using images from Figure 4B and depicting the colocalization between HA-Sumo2 and Shank2 (top panels) and Synapsin 1 (bottom panels). The right panels include the MAP2 (grey) immunolabeling. Scale bar: 5 µm.

**Figure 5—figure supplement 1:**
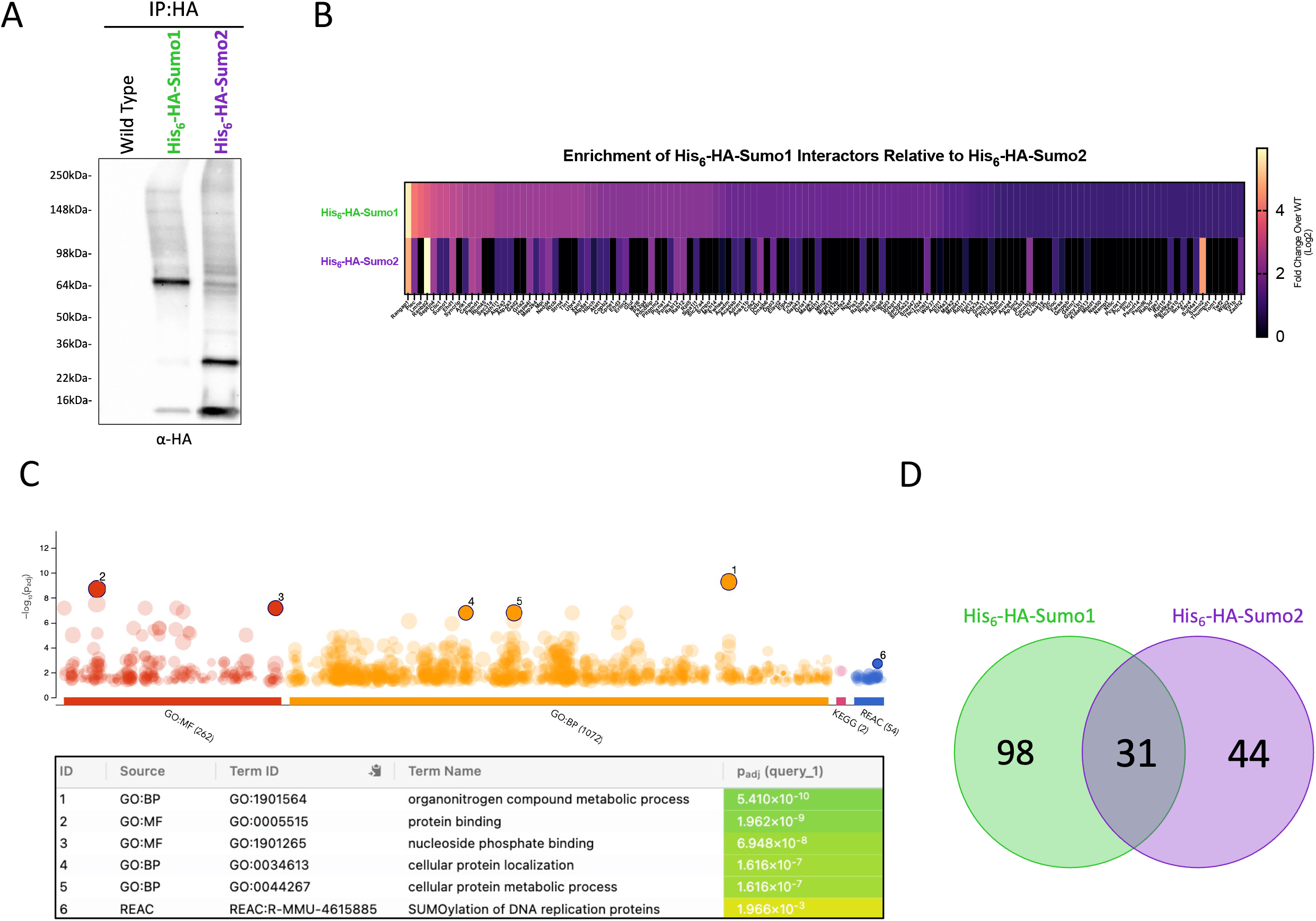
Neuronal Sumo1 has shared and distinct substrates compared to Sumo2 *in vivo*. (A). Anti-HA Western blot analysis of eluates from anti-HA affinity immunoprecipitation (IP: HA) from WT, His_6_-HA-Sumo1, and His_6_-HA-Sumo2 mouse brain lysates demonstrating HA signal patterns of Sumo1 and Sumo2 and their corresponding conjugates (B). Heat map depicting the relative peptide abundance of His_6_-HA-Sumo1 interactors relative to levels in His_6_-HA-Sumo2 immunoprecipitation. (C). gProfiler2 Gene Ontology analysis for His_6_-HA-Sumo1 interactors. (D). Venn diagram depicting unique and common His_6_-HA-Sumo1 and His_6_-HA-Sumo2 interactors.

**Figure 6—figure supplement 1:**
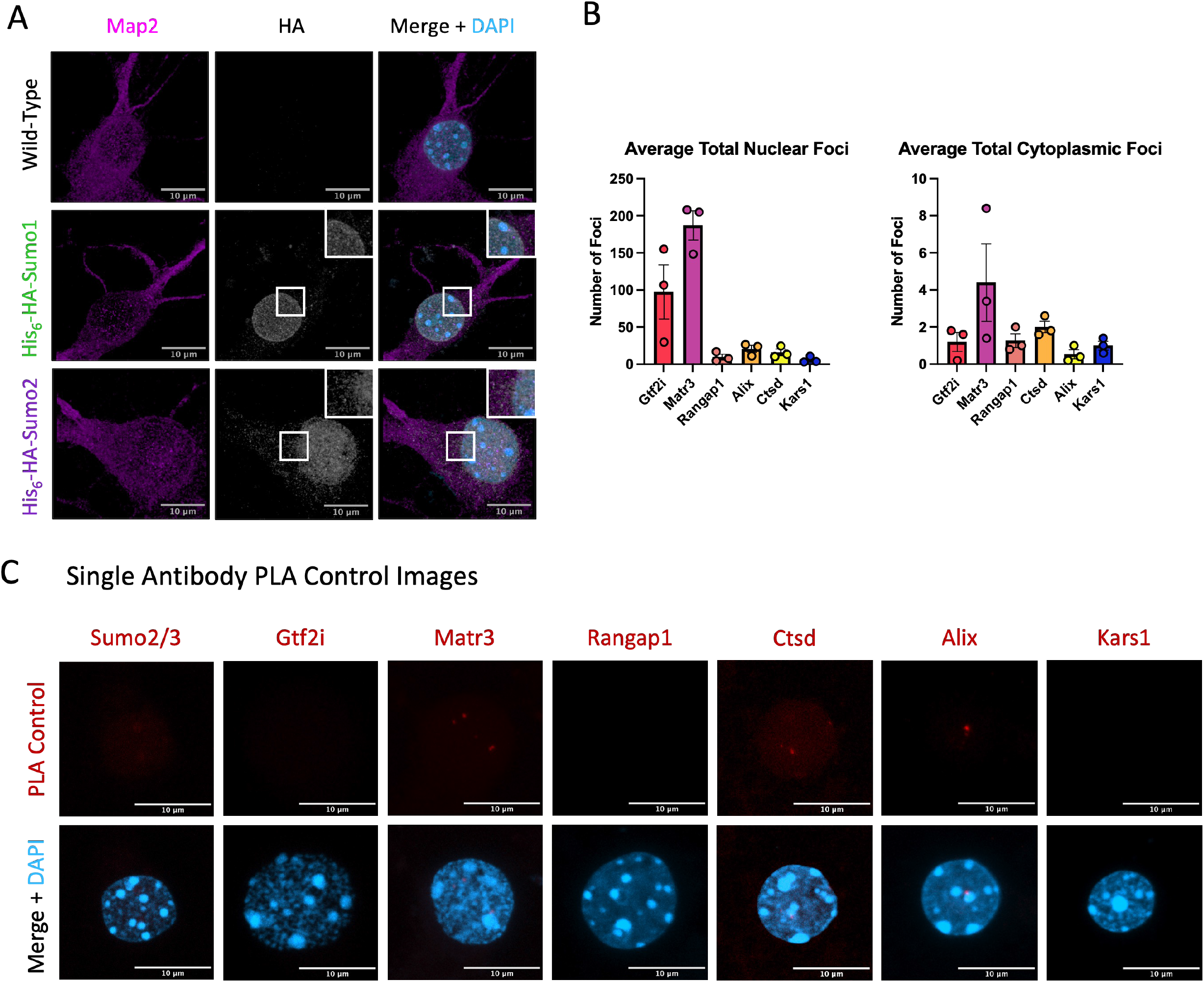
Localization of Sumo2 to extranuclear compartments and proof of PLA assay specificity. (A). Confocal microscopy analysis of anti-Map2 (purple) and anti-HA (grey) immunolabeling of WT (top lanes), His_6_-HA-Sumo1 (middle lanes), and His_6_-HA-Sumo2 primary cortical neurons (bottom lanes). Scale bar: 10 μm. N=3 (B). Bar plot depicting the total number of nuclear (left plot) and cytoplasmic (right plot) PLA foci for the indicated antibody (N=3). Data are presented as a mean ± S.E.M. (C). Representative Z-projected images of proximity ligation assays performed with only a single antibody (indicated on top, top lanes) merged with DAPI (bottom panels). Scale bar: 10 μm.

## ACKNOWLEDGEMENTS

This research was supported in part by an NSERC Discovery Grant and Discovery Launch Supplement to M.W.C.R. (RGPIN-2019-04133 and DGECR-2019-00369); the Canada Research Chairs program to M.W.C.R; Quantitative Synaptology SFB1286/A09 to N.B. and M.T.; the ALS Society of Canada in partnership with the Brain Canada Foundation through the Brain Canada Research Fund, with the financial support of Health Canada, for financial support through the ALS Trainee Award Program 2019 (T.R.S.); an NSERC Undergraduates Student Research Award to T.T.N.; the Ontario Graduate Scholarship (H.M.G.), the Queen Elizabeth II Scholarship (H.M.G.) and a CIHR Canadian Graduate Scholarship (H.M.G.). M.W.C.R. thanks H.Y. Zoghbi (Baylor College of Medicine, HHMI) for initial project discussions, reagent development and the freedom to explore new ideas. The authors also thank all members of the Rousseaux, Brose, and Zoghbi labs for important discussions and critical feedback on the manuscript. The authors also thank the following Core facilities from the University of Ottawa and the Ottawa Hospital Research Institute (OHRI) for use of their facility, equipment, and expertise: the Cell Biology and Imaging Acquisition Core and the OHRI Proteomics Core. The authors also thank the Genome Engineered Rodent Models Core at Baylor College of Medicine and Animal Care and Veterinary Service. Figures 1, 2, 5 and 6 were generated in part with Biorender.com. The views expressed herein do not necessarily represent the views of the Minister of Health or the Government of Canada.

## COMPETING INTERESTS

The authors have no competing interests to declare.

## Notes

### Competing Interest Statement

The authors have declared no competing interest.

